# Disentangling within-cell-type heterogeneity from transcript contamination in imaging-based spatial transcriptomics

**DOI:** 10.64898/2026.11.22.768448

**Authors:** Zhengyang Liu, Yuheng Chen, Can Yang, Yuyao Liu

## Abstract

Recent advances in imaging-based spatial transcriptomics have enabled high-resolution characterization of gene expression and cellular organization in situ. Resolving within-cell-type heterogeneity is essential for understanding how cellular states relate to tissue function and disease. However, imperfect cell segmentation and RNA spillover can contaminate cell-level expression profiles, obscuring intrinsic expression patterns and generating spurious variation. Existing decontamination methods can struggle to distinguish intrinsic variation from contamination when both are inferred from spatial measurements alone, and may suppress genuine state-associated signals when within-cell-type heterogeneity is not explicitly modeled. Here we present Dancer, a reference-guided probabilistic framework that disentangles within-cell-type heterogeneity from transcript contamination by combining cell-type-specific gene programs learned from single-cell data with explicit modeling of intrinsic expression, neighboring-cell contamination and background RNA. Across multiple human tumor datasets with varying gene compositions and panel sizes, Dancer improves cell-type separation and expression specificity while preserving identity-associated signals, achieving superior overall performance in benchmark comparisons. Inferred gene program activities reveal within-cell-type heterogeneity in the form of distinct cellular states and continuous spatial gradients associated with local tissue environments, while reference-guided panel expansion extends molecular characterization beyond the measured genes. Together, Dancer supports accurate transcript decontamination and more faithful interpretation of cellular heterogeneity within its native tissue context.

## Introduction

Understanding how cells are organized within tissues and how they interact is a central goal in biology. Spatial transcriptomics (ST) enables gene expression to be measured within its native tissue context, with imaging-based approaches resolving the identities and positions of individual RNA molecules. Advances in multiplexed RNA imaging, including MERFISH and seqFISH+, have enabled hundreds to thousands of genes to be profiled at subcellular resolution^1, 2^. Platforms such as Xenium and CosMx have extended these capabilities to fixed clinical specimens, supporting detailed maps of cell types and their spatial organization^3, 4^. Recent developments, including Xenium 5K and the Atera whole-transcriptome platform, further expand gene coverage and the range of cellular states that can be resolved in situ^5^. Together, these advances create opportunities to investigate how cells with distinct functional states are organized within tissues and how they interact with the surrounding microenvironment.

Despite these technological advances, transcript contamination remains a major obstacle to interpreting cell-level expression profiles. A common segmentation strategy identifies nuclei from fluorescence staining and expands their boundaries computationally to approximate whole cells. Transcripts detected within each expanded boundary are then aggregated to generate a cell-level expression profile^3, 6^. These approximations can produce segmentation errors that include transcripts from neighboring cells, particularly in densely packed tissues or around cells with irregular shapes^6, 7^. RNA spillover during tissue processing and overlap of cells along the depth of a section can further mix transcripts from different cells, so the measured profiles may not accurately reflect each cell’s own expression^7, 8^. Such contamination can induce apparent co-expression of markers from different lineages and spurious associations between cell state and neighborhood composition, distorting analyses of tissue organization and cell–cell interactions^7, 9^. Both transcript contamination and intrinsic expression can vary across tissue space, making it difficult to determine how much of the observed variation among cells of the same type reflects genuine differences in cellular state. These differences can involve coordinated changes across genes, which can be captured by cell-type-specific gene programs^10^. Separating contamination from intrinsic expression is therefore essential for accurately characterizing within-cell-type heterogeneity and understanding how cellular states are associated with their local microenvironments.

Existing methods have improved the specificity of spatial expression profiles through probabilistic contamination models, neighborhood-based filtering and reference-guided deconvolution^7, 11, 12^. However, disentangling intrinsic within-cell-type heterogeneity from contamination remains challenging. Methods that rely on spatial measurements alone lack an independent expression reference to guide the distinction between intrinsic variation and contamination^7, 11^. Even when a reference is available, correction can suppress genuine within-cell-type heterogeneity if that heterogeneity is not modeled explicitly. For example, anchoring correction to a single expression profile per cell type can lead to genuine state-associated deviations being removed as contamination^12^, weakening coordinated expression signals and obscuring continuous variation among cells of the same type ^13^. A further practical challenge is scalability, as larger cell numbers and broader gene coverage increase the computational demands of spatial analysis^5, 13^.

To address these limitations, we present **Dancer** (**D**isent**AN**gling within-**C**ell-type het**ER**ogeneity from transcript contamination), a reference-guided probabilistic framework for removing contamination while preserving within-cell-type expression variation. Dancer assumes that the single-cell reference is approximately free of transcript contamination and captures the range of intrinsic expression variation present within each cell type in the spatial sample. Based on this assumption, Dancer learns cell-type-specific gene programs from the reference and uses them to represent intrinsic variation in spatial cells. It models observed spatial counts as contributions from intrinsic expression, neighboring-cell contamination and background RNA, jointly inferring each cell’s program activities and source proportions. The inferred intrinsic component yields a decontaminated expression profile for spatial sample, while the estimated program activities and loadings characterize differences in cellular state.

We evaluate Dancer across imaging-based ST datasets from human breast, colorectal and lung cancers spanning different gene panel sizes. Benchmarking against existing methods demonstrates superior overall decontamination performance, with improved cell-type separation and marker specificity. Dancer also uncovers spatially organized variation among cells of the same type. In breast cancer, Dancer resolves B-cell states with distinct receptor-signaling and secretory features, accompanied by differences in memory-associated and cytotoxic expression among neighboring CD8 T cells. In colorectal cancer, Dancer delineates macrophage states and their tumor-associated gradients in program activity while preserving mucosal-defense-associated expression gradients in goblet cells. In lung cancer, Dancer distinguishes mural-like and matrix-associated stromal populations and identifies an alveolar type 2 response state with repair- and transition-associated expression features. Reference-guided panel expansion in breast and lung cancer further extends the molecular characterization of immune and alveolar cell states through imputed expression of genes beyond the measured panels. These results establish Dancer as a framework for disentangling within-cell-type heterogeneity from transcript contamination and interpreting cellular states in their spatial context.

## Results

### Overview of Dancer

Dancer aims to disentangle within-cell-type biological heterogeneity from transcript contamination in imaging-based ST data. It takes as input a single-cell expression reference with cell-type annotations and a spatial dataset comprising a cell-by-gene count matrix, preassigned cell-type labels, and cell coordinates (Fig. 1a). Dancer assumes that the single-cell reference is approximately free of transcript contamination and contains the relevant within-cell-type expression heterogeneity present in the spatial tissue. Additionally, Dancer attributes non-intrinsic transcript to neighboring contamination and background RNA.

**Fig. 1.**
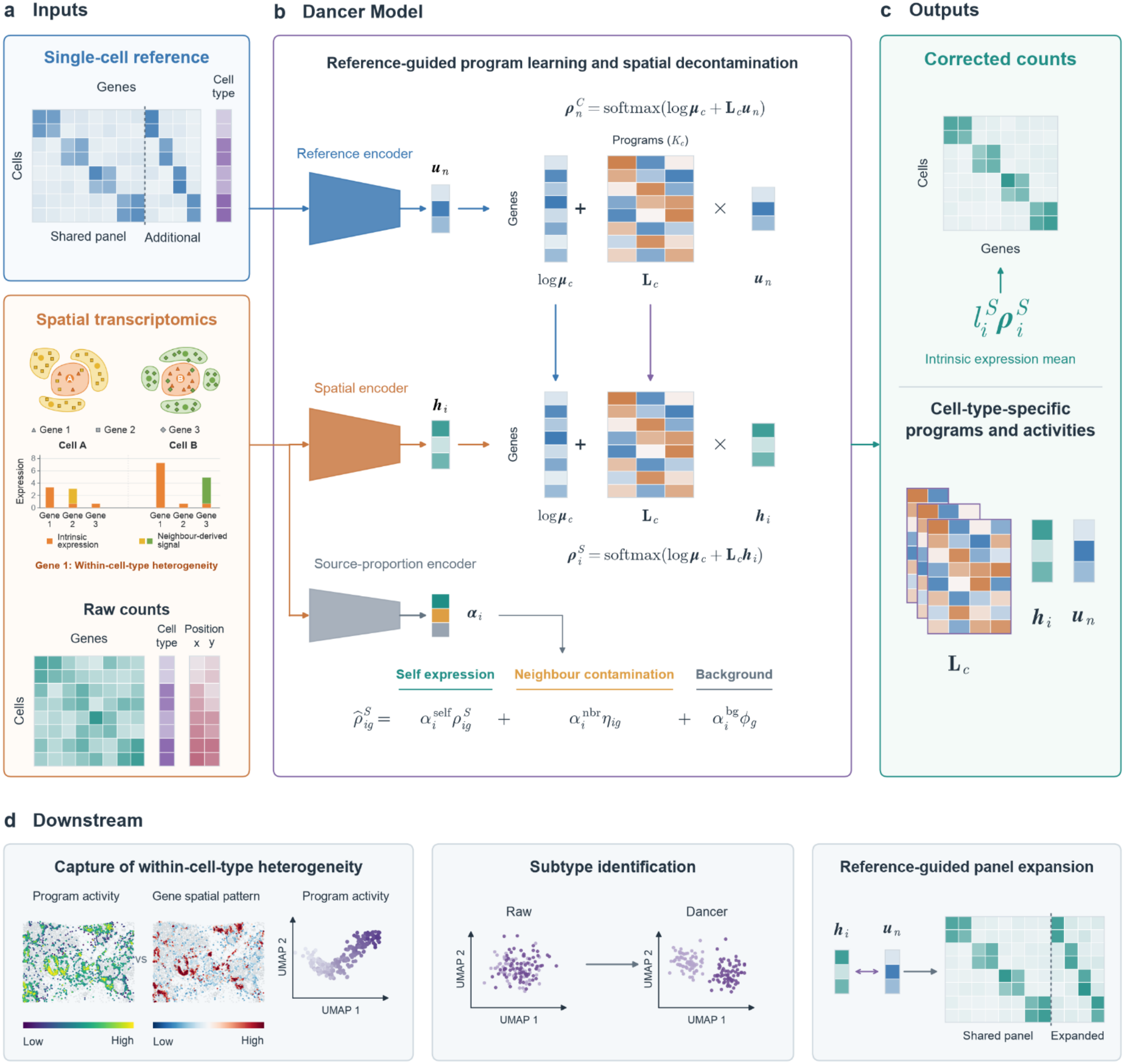
The overview of Dancer. **a, Inputs**. Dancer takes an annotated single-cell reference and a spatial cell-by-gene expression matrix with cell-type labels and coordinates. The schematic illustrates the gene-expression profiles of two cells of the same type in different microenvironments. Differences in Gene 1 reflect intrinsic expression variation, representing within-cell-type heterogeneity, whereas additional signals from Genes 2 and 3 originate from neighboring cells of other types. Such contamination can create apparent heterogeneity if left uncorrected. The reference includes genes shared with the spatial panel and additional genes available for panel expansion. **b, Dancer Model**. A cell-type-specific mean expression profile and reference-learned gene loadings jointly describe intrinsic expression. Separate encoders infer reference and spatial program activities, denoted by *u* and *h*, respectively. Reference programs are learned first, and the fitted gene loadings are then transferred to the spatial model and held fixed during spatial fitting. Vertical arrows indicate that the mean profile and loading matrix are shared from the reference. The spatial expression model combines intrinsic expression, neighbor contamination and background RNA, with cell-specific contributions inferred by a source-proportion encoder. **c, Outputs**. Dancer provides corrected expression profiles, cell-type-specific gene loadings and program activities for individual cells in both modalities. Stacked loading matrices with different widths represent the different numbers of programs selected for different cell types. **d, Downstream**. Dancer’s outputs support spatial characterization of within-cell-type heterogeneity, cellular subtype identification, and reference-guided panel expansion to impute unmeasured gene expression.

To explicitly represent within-cell-type heterogeneity, Dancer models cell-specific variation around a cell-type-specific mean expression profile using gene loadings and program activities. Dancer first fits this model to the single-cell reference. The loadings describe the coordinated expression changes associated with each program, and a reference encoder infers program activities for individual reference cells. Dancer automatically selects the number of programs for each cell type in a data-adaptive manner using the single-cell reference. For spatial cells, Dancer adopts the same intrinsic expression formulation, holding the reference-derived mean profiles and gene loadings fixed while using a separate spatial encoder to infer cell-specific program activities (Fig. 1b). This formulation provides reference-derived biological guidance for separating intrinsic variation from contamination while allowing individual spatial cells to retain their own activity levels.

Dancer models observed spatial counts using a mixture of intrinsic expression, neighboring-cell contamination, and background RNA. A source-proportion encoder estimates the relative contributions of these three sources for each cell (Fig. 1b). The neighborhood component combines the inferred intrinsic expression of nearby cells with spatial proximity and gene-specific spreading tendencies. Background RNA is modeled with a single expression composition shared across the dataset, with a separate contribution estimated for each cell (Methods).

After model fitting, Dancer outputs a decontaminated spatial cell-by-gene expression matrix, cell-type-specific program loadings, and program activities for reference and spatial cells (Fig. 1c). These outputs enable the characterization of within-cell-type heterogeneity across tissue space, the identification of cellular subtypes, and the imputation of the expression of unmeasured genes through reference-guided panel expansion (Fig. 1d). Dancer thus links cellular states to their local tissue environments, supporting a more faithful and comprehensive interpretation of this heterogeneity.

### Dancer outperforms existing methods in accurate transcript decontamination

To evaluate decontamination performance, we compared Dancer with four baseline methods, ResolVI^7^, DenoIST^11^, MisTIC^14^ and SPLIT^12^, alongside uncorrected expression (Raw). Comparisons were conducted across multiple imaging-based ST datasets, with preprocessing, baseline configurations and evaluation procedures described in Methods.

Transcript contamination from neighboring cells and background RNA can obscure cell-type-specific expression patterns, reducing separability between cell types. We therefore assessed cell-type separability using average silhouette width (ASW), with higher scores indicating clearer separation. Dancer increased ASW from 0.512 to 0.571 in breast cancer and from 0.515 to 0.611 in colorectal cancer, achieving the largest improvement over Raw among all evaluated methods (Fig. 2b, c). Consistently, the breast cancer UMAP showed substantial overlap between several populations in Raw, whereas Dancer resolved more distinct cell-type-associated groups, making differences in their expression profiles more apparent (Fig. 2a).

**Fig. 2.**
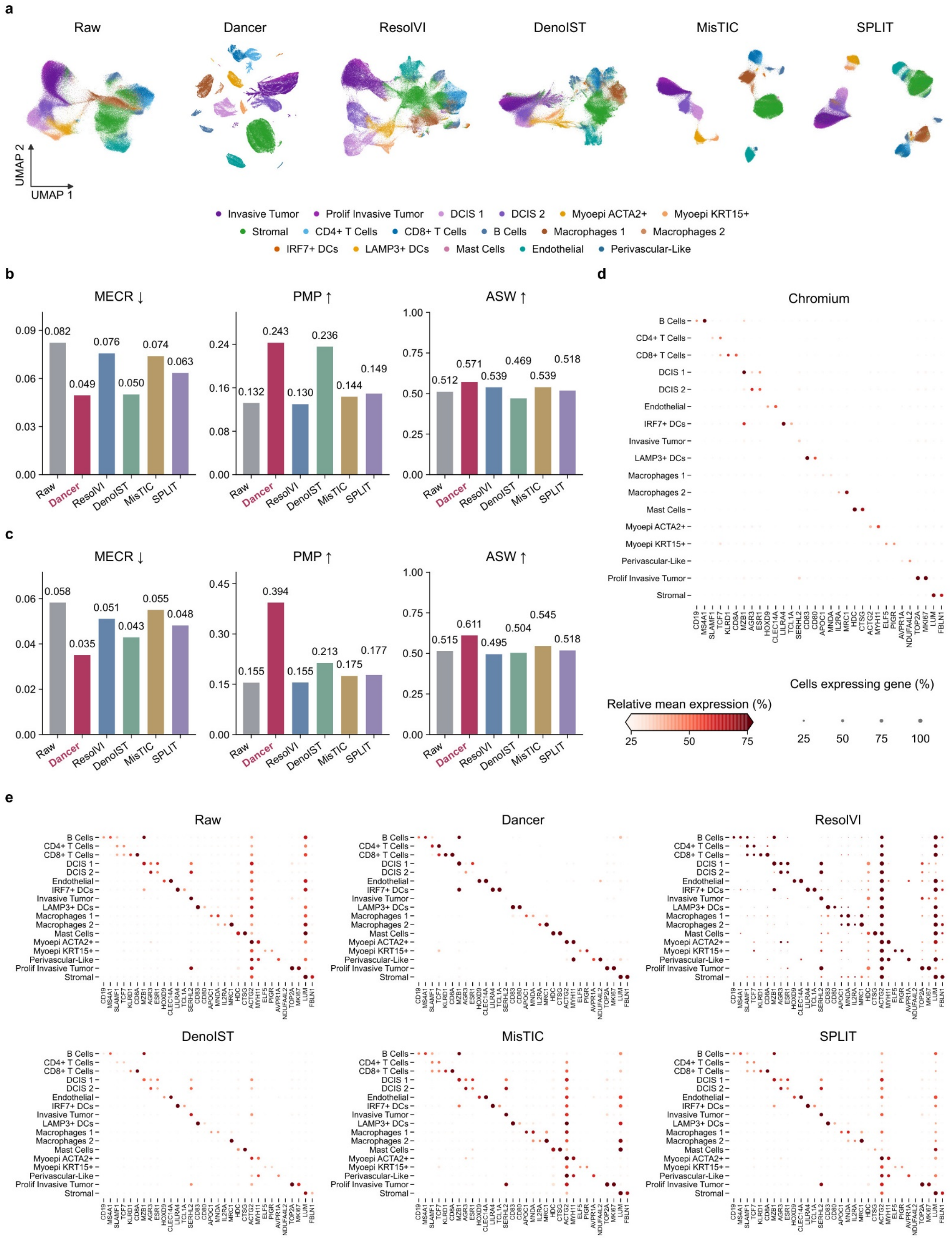
Dancer improves transcript decontamination and preserves cell-type-associated expression. **a**, UMAP visualization of breast cancer expression profiles from Raw, Dancer, ResolVI, DenoIST, MisTIC and SPLIT, colored by the same cell-type annotations. Cells with at least 20 total counts are displayed. **b, c**, mutually exclusive co-expression rate (MECR), positive marker purity (PMP) and average silhouette width (ASW) in breast cancer (**b**) and colorectal cancer (**c**). Lower MECR and higher PMP and ASW indicate better performance. ASW is rescaled to 0–1 and calculated on the common set of cells with at least 20 total counts in every compared expression matrix; MECR and PMP are calculated using all input cells. **d, e**, expression of the same 32 representative markers across 17 cell types in the breast cancer dataset, shown for the Chromium single-cell reference (**d**) and Raw and corrected spatial profiles (**e**). Dot size represents the percentage of cells with count-scale expression ≥0.5, while color represents mean log-normalized expression, scaled for each gene by its maximum across all displayed methods and cell types. Expression is normalized to 10,000 counts per cell before log transformation.

Effective decontamination should reduce ectopic marker expression while preserving signals associated with cell identity. We evaluated this balance using the mutually exclusive co-expression rate (MECR), which measures co-expression between markers of different cell types, and positive marker purity (PMP), which measures the proportion of expression contributed by a cell’s own-type markers. Compared with Raw, Dancer reduced MECR from 0.082 to 0.049 and increased PMP from 0.132 to 0.243 in the breast cancer dataset (Fig. 2b). Although DenoIST achieved marker-based scores comparable to Dancer’s on this dataset, several cell-type markers showed reduced expression in its corrected profiles (Fig. 2e), consistent with its filtering procedure, which either retains each cell–gene count in full or sets it to zero^11^. Dancer’s advantage was more pronounced in colorectal cancer, where it reduced MECR from 0.058 to 0.035 and increased PMP from 0.155 to 0.394, attaining the lowest MECR and highest PMP among all evaluated methods (Fig. 2c). Together with the marker profiles, these results support the removal of ectopic expression alongside preservation of cell-type-associated signals.

Representative marker dot plots also provided direct gene-level evidence of improved expression specificity and preservation of cell identity (Fig. 2d, e). The plots display 32 representative markers across 17 cell types in breast cancer, with markers ordered by their associated cell types to form diagonal blocks. We examined whether corrected profiles more closely followed the Chromium-supported expression pattern, with stronger signals concentrated in these blocks. Dancer reduced ectopic marker expression while preserving expression in the expected cell types. For example, ACTG2 was broadly detected across cell populations in the raw spatial data. Among the methods evaluated, Dancer most closely matched the ACTG2 expression pattern observed in the Chromium single-cell reference, with expression largely restricted to ACTA2+ myoepithelial cells (Fig. 2d, e).

Together, these results show that Dancer improves cell-type separability and expression specificity while preserving identity-associated signals.

### Dancer improves expression fidelity and uncovers meaningful cellular heterogeneity in the breast cancer immune microenvironment

Reliable interpretation of the tumor immune microenvironment depends on assigning expression signals to the appropriate cells and tissue compartments. In the breast cancer dataset, we first evaluated Dancer’s ability to improve expression fidelity by testing how it affected cell-cell communication and spatial expression (Fig. 3a)^3^. CellChat^15^ analysis showed that communication networks inferred from Dancer-corrected profiles agreed more closely with the matched Chromium reference than those inferred from Raw, with edge precision increasing from 0.391 to 0.718 while recall also improved slightly (Fig. 3b). For PTPRC–MRC1, correction removed spurious connections inferred from Raw and recovered the same set of sender–receiver combinations as the reference. Moreover, after Dancer correction, ACTA2 expression was more concentrated in the myoepithelial layer of a duct identified in the matched H&E image^3^, with reduced signal outside that layer (Fig. 3c). Accordingly, the ability of ACTA2 expression to distinguish these populations increased from an AUROC of 0.923 to 0.996, and its Spearman correlation with the myoepithelial label increased from 0.478 to 0.926 (Fig. 3c, Methods). These complementary observations linked improved expression fidelity to both inferred cellular communication and anatomical organization.

**Fig. 3.**
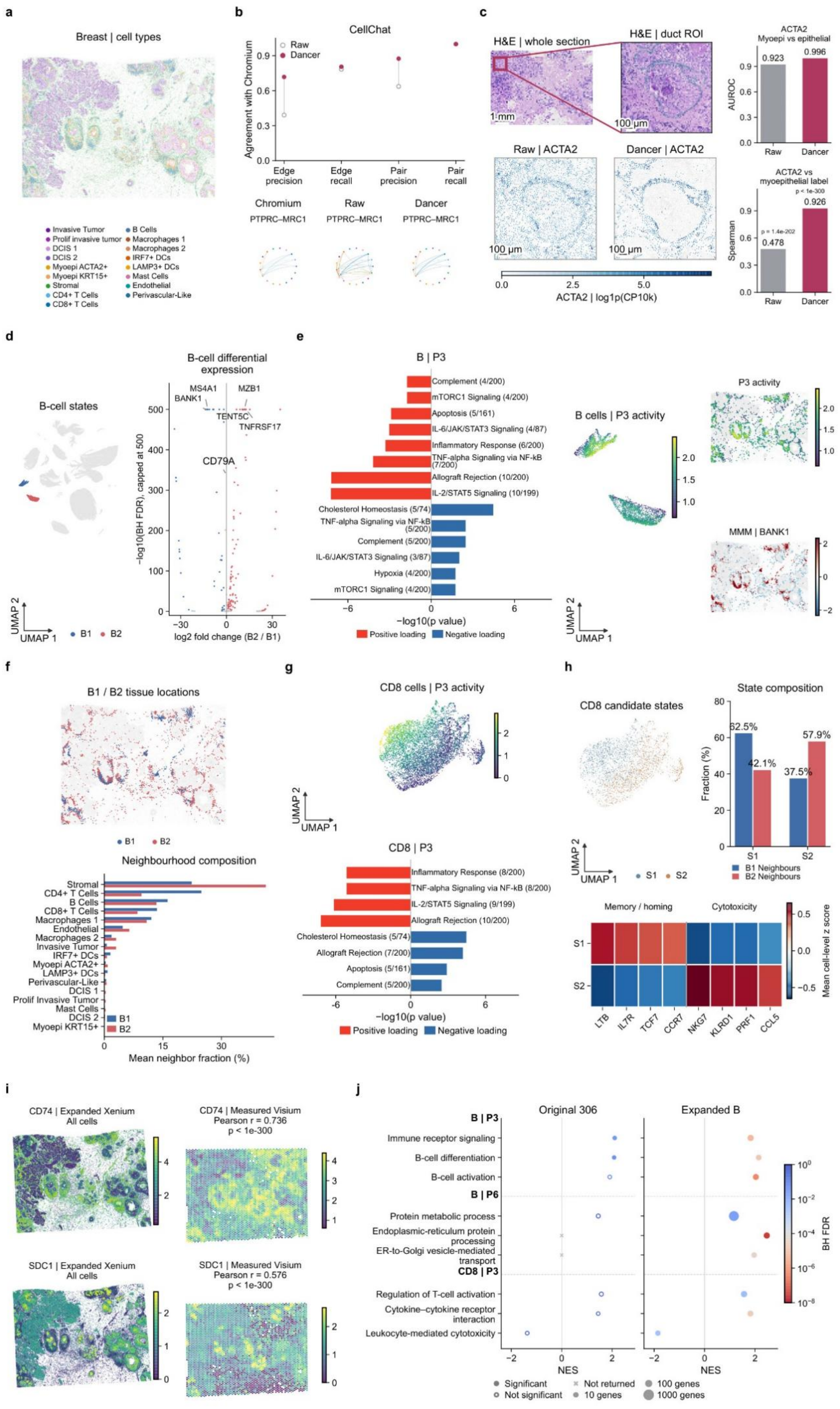
Dancer improves expression fidelity and reveals spatially organized immune heterogeneity in breast cancer. **a**, spatial distribution of 158,030 Xenium cells colored by their 17 annotated cell types. **b**, concordance of CellChat-inferred communication with the matched Chromium reference, evaluated by precision and recall for sender–receiver–ligand–receptor edges and for unique ligand–receptor pairs. Open and filled points indicate Raw and Dancer, respectively. PTPRC–MRC1 networks show significant connections (*P* < 0.05); nodes represent cell types, arrows indicate sender-to-receiver direction, and line widths indicate communication probabilities on a shared scale. Node colors follow **a. c**, whole-section and enlarged duct H&E images from the matched breast cancer dataset^3^, registered to Xenium coordinates, with Raw and Dancer-cleaned ACTA2 expression in the same region. Green overlays identify annotated myoepithelial cells. Expression maps share a log1p(CP10k) color scale. AUROC measures separation of 361 myoepithelial cells from 3,203 adjacent epithelial or tumor cells by ACTA2 expression; Spearman correlation measures its association with the binary myoepithelial label. Scale bars, 1 mm in the whole-section image and 100 μm in the enlarged views. **d**, B1 and B2 in the Dancer whole-section expression UMAP and their differential-expression volcano plot. B1/B2 denote the two groups identified in this analysis. The volcano plot shows log2 fold change (B2/B1) against −log10 Benjamini–Hochberg-adjusted *P* from two-sided Wilcoxon rank-sum tests, with the vertical display capped at 500. **e**, hallmark enrichment of B-cell P3, its activity in UMAP and tissue space, and the BANK1 spatial pattern estimated by MMM. Enrichment uses the 100 highest- and 100 lowest-loading genes, tested separately by one-sided Fisher’s exact tests with the Enrichr default background; displayed terms have Benjamini–Hochberg-adjusted *P* ≤ 0.05. Red and blue bars denote the high- and low-loading ends, respectively, and bar length represents −log10 unadjusted *P*; labels give the number of overlapping genes relative to the full gene-set size. P3 activity correlated with the MMM-estimated BANK1 pattern across 4,862 B cells (Spearman *r* = 0.329, two-sided *P* = 7.72 × 10^−123^). **f**, B1/B2 tissue locations and mean cell-type proportions among neighbors within 30 μm of each B cell. **g**, CD8 T-cell P3 activity on a CD8-only expression UMAP and Hallmark enrichment calculated as in e. Program indices are specific to each cell type. **h**, CD8 T-cell candidate states defined using P3/P4 activity, displayed on the expression UMAP; their proportions among CD8 T cells exclusively neighboring B1 or B2; and state-averaged, gene-wise standardized expression of memory/homing- and cytotoxicity-associated genes. **i**, predicted CD74 and SDC1 expression in all Xenium cells after panel expansion and corresponding measured Visium expression. Pearson correlations are calculated across 3,922 matched spots after aggregating predicted cell counts and normalizing each spot. Correlation *P* values in **c, e** and **i** are two-sided. **j**, functional enrichment associated with B-cell P3/P6 and CD8 T-cell P3 in the original 306-gene and expanded panels. Genes are ranked by Spearman correlation between expression and program activity for preranked GSEA. Position indicates normalized enrichment score (NES), color indicates Benjamini–Hochberg-adjusted *P*, and size indicates the number of leading-edge genes. Filled points meet the display threshold (adjusted *P* ≤ 0.05 for GO Biological Process and ≤ 0.10 for KEGG); open points do not, and crosses indicate terms without a returned enrichment result.

Alongside these improvements, the corrected profiles retained heterogeneity within the B-cell compartment, which separated into two groups in the expression UMAP, designated B1 and B2 (Fig. 3d, Methods). B1 expressed higher levels of BANK1 and MS4A1, whereas B2 was enriched for MZB1, TENT5C and PRDM1, suggesting a distinction between B-cell receptor-associated identity and a secretory, plasma-cell-like expression state. This interpretation is consistent with the role of BANK1 in B-cell receptor signaling^16^ and the involvement of MZB1 and TENT5C in immunoglobulin processing and secretion^17, 18^. Consistent with this distinction, B-cell P3 program activity inferred by Dancer was higher in B1 and lower in B2, and its tissue distribution corresponded to the BANK1 spatial pattern estimated separately using the MMM^19^. Hallmark over-representation analysis^20^ of the highest- and lowest-loading genes further associated the high-loading end of P3 with IL-2/STAT5, TNF–NFκB and inflammatory-response signatures, while the low-loading end included cholesterol-homeostasis and metabolic signatures (Fig. 3e). Together with the differential expression of BANK1 and MS4A1 versus MZB1 and TENT5C, these enrichment patterns supported a distinction between receptor- and immune-associated expression in B1 and a secretory state with accompanying metabolic features in B2.

Because local B–T-cell organization can accompany distinct immune activities in breast cancer^21^, we next examined whether the two B-cell states occupied different cellular neighborhoods. The local cellular composition differed between the two states, with higher proportions of both CD4 and CD8 T cells around B1 than around B2 (Fig. 3f). Thus, the B-cell expression states were associated with differences in the composition of the local immune microenvironment, motivating a closer examination of the functional states of neighboring T cells.

We next examined whether the expression states of CD8 T cells were associated with the state of neighboring B cells (B1 or B2). The P3 program activity of CD8 T cells formed a continuous gradient across an independently computed expression UMAP, with its high-loading genes enriched for IL-2/STAT5 and TNF–NFκB signaling and inflammatory-response signatures (Fig. 3g). Clustering of CD8 T cells identified two candidate states, which we designated S1 and S2 (Fig. 3h, Methods). S1 showed higher P3 activity and higher expression of LTB, IL7R, TCF7, and CCR7, consistent with memory- and homing-associated features^22^. S2 showed higher expression of NKG7, KLRD1, PRF1, and CCL5, consistent with a cytotoxic effector-like expression profile. Among CD8 T cells exclusively neighboring B1 cells, 62.5% were in S1. Among those exclusively neighboring B2 cells, 57.9% were in S2 (Fig. 3h). The enrichment of P3-high CD8 cells around B1 was consistent with the immune-signaling features observed in B1, linking the local B-cell state to the functional composition of nearby T cells.

Finally, we used Dancer’s program-activity representation of spatial and reference cells of the same type to extend the molecular interpretation beyond the measured panel. Drawing on the approximately contamination-free reference, we predicted unmeasured gene expression for each spatial cell by averaging the expression profiles of several same-type reference cells with the most similar program activities. We examined CD74 and SDC1 as additional readouts of antigen-presentation-related and plasma-cell-associated expression, respectively. Within the B-cell compartment, predicted CD74 expression was higher in B1 and positively associated with P3 activity, whereas SDC1 was higher in B2 and negatively associated with P3. Their whole-section maps additionally revealed distinct spatial distributions, which agreed with matched Visium measurements, with spot-level Pearson correlations of 0.736 for CD74 and 0.576 for SDC1 (Fig. 3i, Methods). Gene set enrichment analysis based on expression–activity correlations^23^ further strengthened the functional interpretation (Fig. 3j): immune receptor signaling and B-cell differentiation associated with B-cell P3 gained stronger statistical support, and B-cell activation became significant. For the P6 program of B cells, whose activity was higher in B2, the expanded panel supported protein metabolism and additionally revealed enrichment of endoplasmic-reticulum protein processing and ER-to-Golgi transport, reinforcing the secretory interpretation of this state. The P3 program likewise gained support for T-cell activation and cytokine signaling at its positive expression–activity end and cytotoxicity at its negative end.

These results show that Dancer improves expression fidelity and resolves within-cell-type heterogeneity in both B cells and CD8 T cells, revealing spatial associations between their distinct states in the breast cancer immune microenvironment.

### Dancer delineates cell subtypes while preserving tumor-associated spatial gradients in colorectal cancer

To examine expression fidelity within the spatially structured colorectal tumor, we first assessed expression mixing between cells and their neighbors from different cellular lineages (Fig. 4a, b, Methods). Dancer reduced this cross-lineage similarity across all six broad compartments, while increasing cell-type expression agreement with Chromium from 0.687 to 0.766, outperforming baseline methods. We also examined the spatial distribution of ACTA2, which encodes α-smooth muscle actin and is associated with contractile stromal populations, including myofibroblasts located around colonic crypts and within tumor-associated stroma^24^. Compared with the diffuse signal in Raw, corrected ACTA2 expression was more spatially restricted and agreed more closely with matched Visium HD and Visium measurements from the same specimen (Fig. 4c, Methods)^25^. These results linked reduced expression mixing between distinct cell populations to improved agreement with reference profiles and tissue-level expression patterns.

**Fig. 4.**
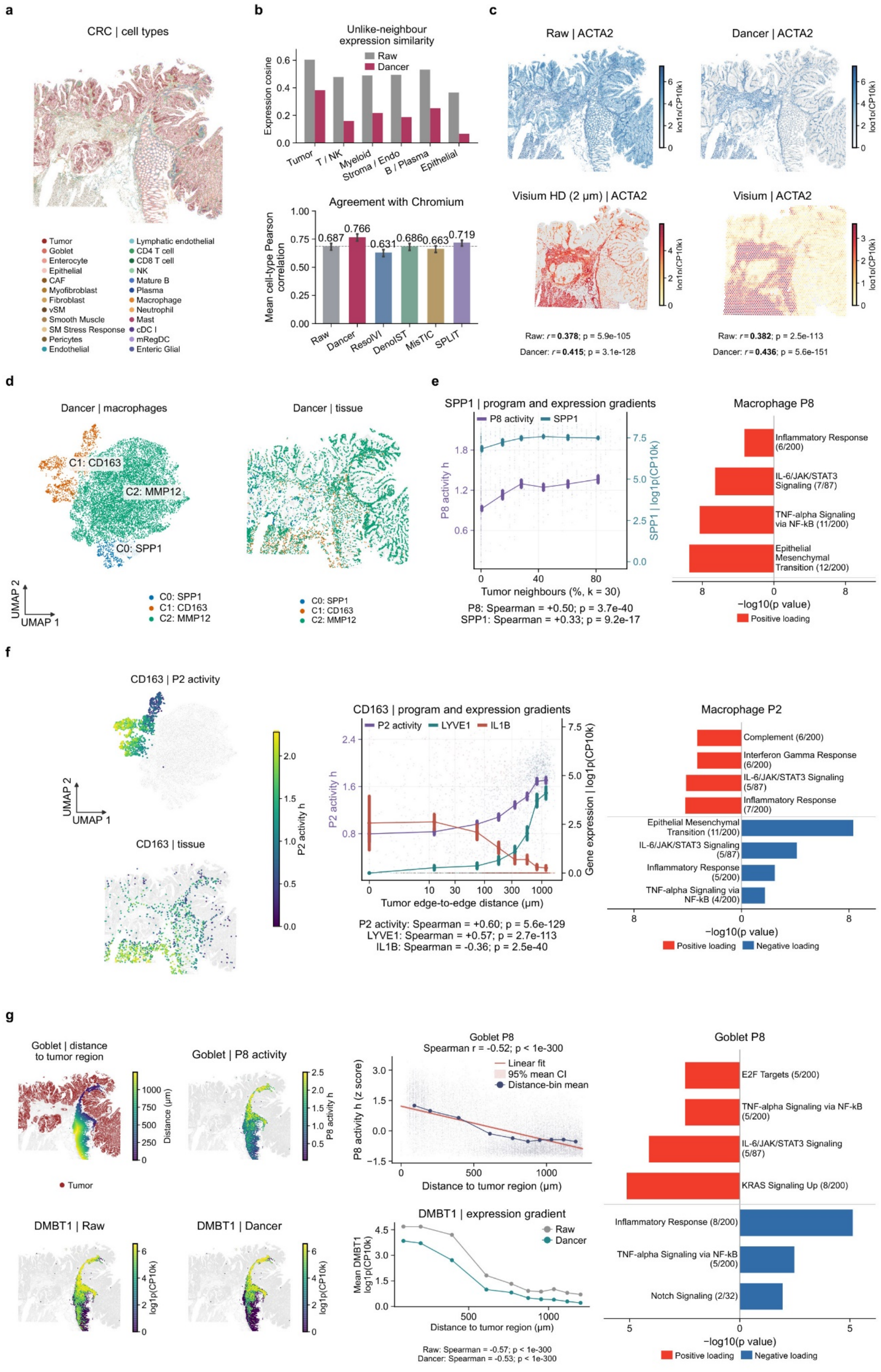
Dancer delineates macrophage states and preserves tumor-associated spatial gradients in colorectal cancer. **a**, spatial distribution of annotated cell types in the colorectal cancer Xenium section. **b**, Top, mean cosine similarity between each cell’s expression profile and the aggregated profile of neighbors from different cellular lineages, selected among its 15 nearest neighbors and summarized by six broad focal-cell compartments. Bottom, agreement with the Chromium reference was quantified by calculating a Pearson correlation across the 407 shared genes for each of the 23 shared cell types and averaging the correlations equally across cell types. Normalized expression was averaged within each cell type and then rescaled for each gene to sum to one across cell types. Error bars indicate 95% confidence intervals from 400 bootstrap resamples of genes. **c**, ACTA2 expression in Raw and Dancer Xenium profiles, as well as in the Visium HD and Visium samples from the matched colorectal cancer specimen^25^. Maps show log-normalized expression, with a common color scale for Raw and Dancer. The reported Pearson correlations compare each Xenium profile with the corresponding external platform after aggregation into registered 100 × 100 μm grids and normalization over the 407 shared genes. Raw and Dancer use the same 3,072 grids for Visium HD and 3,243 grids for Visium. **d**, macrophage-only expression UMAP and tissue locations for 12,000 sampled macrophages. Five initial Leiden clusters were consolidated by marker expression into SPP1, CD163 and MMP12 states; other tissue cells are gray. **e**, relationships of P8 program activity and SPP1 expression with the percentage of tumor cells among the 30 nearest neighbors of each SPP1-state macrophage, and Hallmark enrichment of the high-loading end of macrophage P8 program. **f**, P2 activity in CD163-state macrophages in UMAP and tissue space, P2 activity and LYVE1/IL1B expression versus tumor-edge distance, and Hallmark enrichment of both loading ends of macrophage P2. Distance is the nearest tumor-cell centroid distance minus the two cells’ area-equivalent radii, truncated at zero. In **e** and **f**, individual points represent cells and lines connect binned means with error bars of ±1.96 standard errors; the left axis shows program activity and the right axis shows log1p(CP10k) gene expression. **g**, spatial maps of goblet-cell distance to tumor regions and P8 activity, the association between P8 activity and tumor distance, Raw and Dancer DMBT1 expression maps and distance profiles, and Hallmark enrichment at both ends of the goblet-cell P8 loading distribution. In the distance map, goblet cells are colored by distance to the nearest tumor region, tumor cells are shown in red, and all remaining cells are shown in gray. Cells inside a tumor region were assigned a distance of zero. Distance-trend analyses included 17,799 goblet cells at or below the 95th percentile of distance to the density-defined tumor regions. The activity–distance plot shows standardized P8 activity, a linear fit with a 95% confidence band for the mean, and distance-bin means. The Raw and Dancer DMBT1 maps share a color scale, and the curves show mean log1p(CP10k) expression in the same distance bins. Spearman correlations in **e–g** were calculated across individual cells. Correlation tests in **c** and **e–g** were two-sided.

We next focused on macrophages, which encompass functionally diverse populations occupying distinct tissue environments in colorectal cancer^26, 27^. Clustering of macrophage expression profiles, followed by marker-based annotation, identified three states characterized by SPP1, CD163 and MMP12 expression (Fig. 4d). These states also differed in their tissue distributions: MMP12 macrophages were broadly distributed, SPP1 macrophages showed localized accumulations along parts of the left side of the section, and CD163 macrophages were prominent in the lower part of the section, linking their expression differences to distinct spatial arrangements.

Beyond the differences between these states, we next examined whether program activity captured continuous variation within each population in relation to the local tumor environment. SPP1+ macrophages have been reported to accumulate in colorectal tumors and exhibit extracellular-matrix- and tissue-remodeling-associated expression^28^. Within the SPP1 state, P8 program activity varied across cells and increased with the fraction of tumor cells among nearby neighbors, alongside an increase in gene SPP1 expression (Spearman correlations of 0.50 and 0.33, respectively; Fig. 4e, Methods). Hallmark enrichment linked the highest-loading genes of P8 to TNF–NFκB and IL-6/JAK/STAT3 signaling, inflammatory responses and the epithelial–mesenchymal transition gene set, with SPP1, TGFBI and THBS1 in the latter supporting a tissue-remodeling interpretation.

Within the CD163 population, P2 program activity showed a clear gradient across the expression UMAP and tissue space (Fig. 4f). P2 activity increased with distance from the tumor edge (Spearman correlation, 0.60), accompanied by increasing LYVE1 expression and decreasing IL1B expression. Its high-loading genes included the complement-associated C1QA and C1QC and the scavenging-associated CD163 and MRC1, whereas IL1B and IL6 contributed to inflammatory signatures at the low-loading end. Together, these findings suggest a shift from IL1B-associated inflammatory expression near tumors toward resident-like and complement/scavenging-associated expression at more distant sites, consistent with the spatial segregation of macrophage states reported previously^27^.

We next examined whether tumor-associated spatial variation also extended to the epithelial compartment. The P8 program activity of goblet cell was higher near tumor regions and decreased with distance (Spearman correlation, −0.52; Fig. 4g, Methods), with its highest-loading genes enriched for IL-6/JAK/STAT3 and E2F targets, suggesting cytokine-response and cell-cycle-related variation. DMBT1, a positively loaded gene whose expression correlated with P8 activity, provided a complementary link to epithelial defense through its established role in intestinal mucosal protection^29^. Its expression was higher near tumor regions and declined with increasing distance, a gradient retained after Dancer correction (Spearman correlations of −0.57 and −0.53 in Raw and Dancer, respectively).

Taken together, these results show that Dancer improves expression specificity and cross-platform agreement while resolving macrophage states and preserving tumor-associated gradients consistent with myeloid functional diversity and epithelial mucosal defense.

### Dancer scales to large panel and identifies a repair-associated AT2 cell state in lung cancer

We next applied Dancer to a lung cancer section profiled with Xenium Prime 5K, extending the analysis to a larger panel with 4,598 genes shared with the single-cell reference (Fig. 5a). Dancer improved cell-type separation in the expression UMAP, with higher ASW and PMP and lower MECR than Raw (Fig. 5b). Using paired Wilcoxon signed-rank tests on fixed gene-pair sets defined by the scVI-purified reference, we found that correction strengthened reference-supported co-expression within lineages while reducing putatively artifactual co-expression across lineages (Fig. 5c, Methods). We further examined expression differences between stromal cells near and far from tumors. Tumor-associated genes, including EPCAM, MUC1 and MAL2, showed attenuated near–far contrasts after correction, whereas stromal genes such as THBS2 and POSTN retained their higher expression near tumors (Fig. 5d). These observations suggested that removal of ectopic tumor-associated expression preserved spatially associated differences within the stromal compartment.

**Fig. 5.**
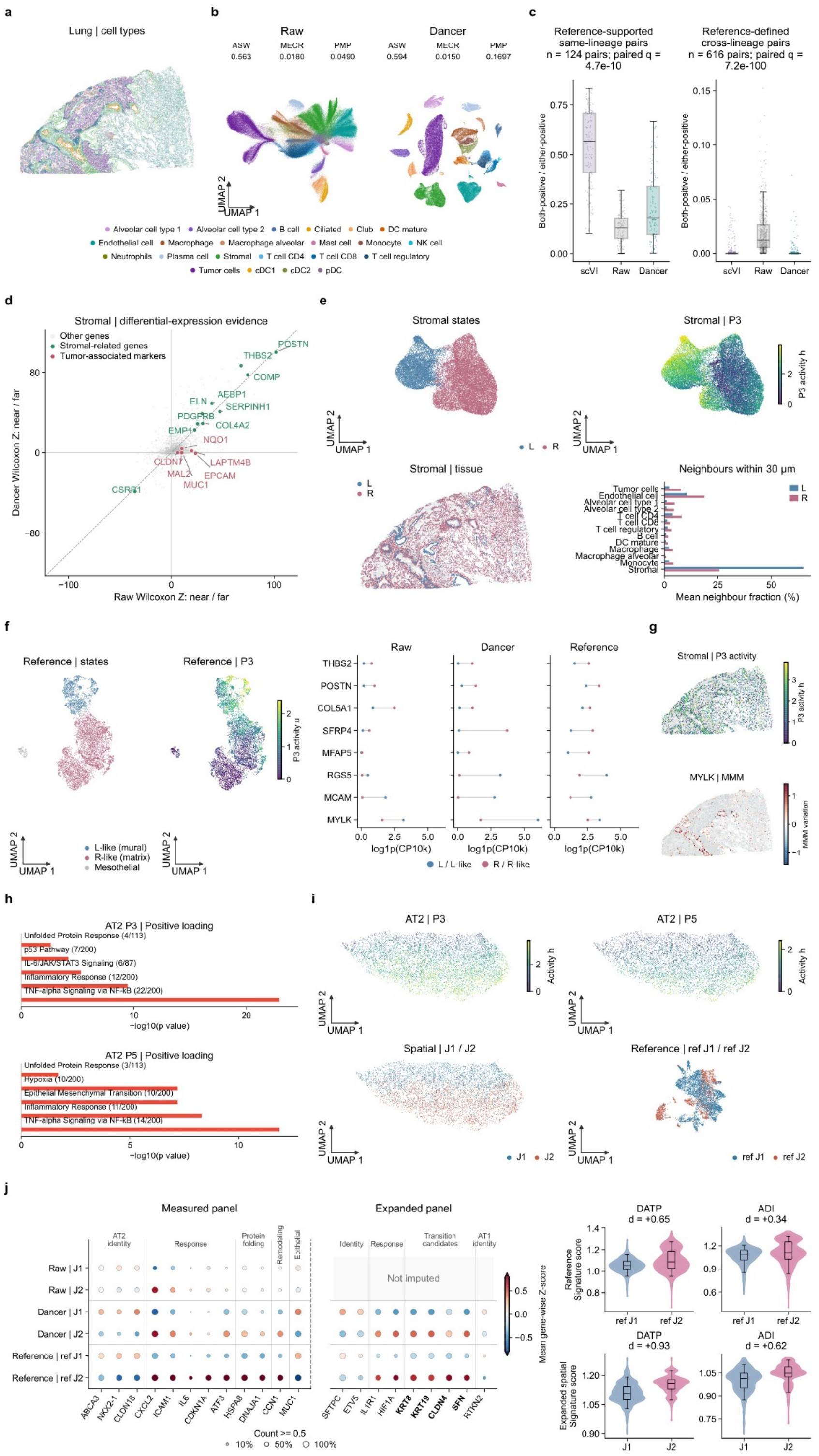
Dancer restores cell-intrinsic expression and reveals stromal and alveolar cell-state heterogeneity in lung cancer. All reference panels use the LuCA single-cell reference processed with scVI, which also served as the reference for Dancer model fitting. **a**, spatial distribution of annotated cell types in the lung cancer Xenium section. **b**, Raw and Dancer expression UMAPs, colored by the annotations in **a** and displaying cells with total expression ≥20. ASW, MECR and PMP are calculated as defined in Methods, with ASW evaluated on the shared eligible cell set. **c**, gene-pair co-expression in the scVI-processed reference, Raw and Dancer, measured as the fraction of cells positive for both genes among those positive for at least one. The reference defined 124 same-lineage pairs (co-expression ≥ 0.10) and 616 cross-lineage pairs (≤ 0.05). Each point represents a gene pair; boxes show medians and interquartile ranges, and whiskers extend to the most extreme observations within 1.5 interquartile ranges. Raw–Dancer comparisons use two-sided paired Wilcoxon signed-rank tests, with Benjamini–Hochberg correction across the four co-expression and reference-deviation comparisons described in Methods. **d**, signed Wilcoxon rank-sum Z statistics comparing the same stromal cells near (≤20 μm) and far (≥80 μm) from tumor cells, calculated separately for Raw and Dancer. Distance is measured between cell edges using area-equivalent radii; positive Z indicates higher expression near tumors. Stromal-associated genes are shown in green, tumor-associated markers in pink, and other genes in gray. The dashed diagonal marks equal Wilcoxon Z statistics for Raw and Dancer. **e**, expression-derived stromal L/R states, P3 activity on the same UMAP, their tissue distributions and mean cell-type fractions among neighbors within 30 μm. Other cells are shown in gray in the spatial map. **f**, independently clustered reference stromal cells, reference P3 activity and state-averaged expression of eight identity-associated genes in Raw, Dancer and the reference. Reference groups are matched to spatial L/R states using gene expression and the original LuCA annotations^31^; the small mesothelial group is shown separately in gray. Points in the expression panels show mean log1p(CP10k) expression, with blue indicating L/L-like and pink indicating R/R-like cells. **g**, spatial P3 activity and the MYLK pattern estimated using MMM, shown for the same 4,000 stromal cells. P3 activity correlated with the MMM-estimated MYLK pattern (Spearman *r* = 0.300, two-sided *P* = 3.31 × 10^−84^), supporting the spatial variation in MYLK (*q* = 6.50 × 10^−104^). **h**, selected Hallmark enrichments among the 100 highest-loading genes of AT2 P3 and P5, tested using one-sided Fisher’s exact tests with the Enrichr default background. Displayed terms have Benjamini–Hochberg-adjusted *P* ≤ 0.05; bar lengths show −log10 unadjusted *P*, and labels give the overlapping-gene count relative to the full gene-set size. **i**, AT2 expression UMAPs show P3 and P5 activities and J1/J2 states in spatial cells, as well as ref J1/ref J2 states in the scVI-processed reference. J2 and ref J2 denote the groups with higher mean standardized activity across the two programs. **j**, state-associated expression within the measured and expanded panels, as well as the DATP and ADI signature scores in reference and expanded spatial cells. Dot color indicates the mean gene-wise Z-score within each state, calculated by standardizing log-normalized expression separately within each modality. Dot size indicates the fraction of cells with count-scale expression ≥ 0.5. Spatial J1/J2 assignments are held fixed across Raw, Dancer and panel-expanded profiles. The blank area in the expanded panel indicates unavailable Raw predictions. Bold gene labels highlight literature-supported transitional-state markers^33, 34^. Signature scores use all represented genes from the published human DATP set^35^ and ADI set^33^, comprising 147 and 353 genes, respectively. Scores are calculated as mean log-normalized signature expression minus the mean of expression-matched control genes. Violin plots show cell-level score distributions, with inset boxes indicating medians and interquartile ranges; Cohen’s *d* summarizes J2−J1 differences within each modality. Program indices are cell-type-specific.

Then, we further examined the heterogeneity within stromal cells, which formed two expression clusters on the left and right of the UMAP, designated L and R (Fig. 5e, Methods). P3 program activity was higher in L and lower in R, with Hallmark enrichment linking its high-loading end to myogenesis and its low-loading end to extracellular-matrix-related genes within the epithelial–mesenchymal transition signature, including MFAP5 and SFRP4. Higher RGS5, MCAM and MYLK expression in L supported a mural, vascular-wall-like identity, whereas SFRP4 and MFAP5 favored a matrix-associated interpretation of R, consistent with the diversity of mesenchymal populations in the human lung^30^. The spatial distribution of P3 also corresponded to the pattern estimated by MMM for the contractility-related gene MYLK (Fig. 5g). These populations occupied different local environments, with higher fractions of tumor, endothelial and immune cells around R (Fig. 5e), linking their expression identities to distinct tissue contexts.

To assess whether these spatial populations had counterparts in the single-cell reference, we independently clustered reference stromal expression profiles and identified two principal groups with corresponding P3 activity differences (Fig. 5f). The P3-high, L-like group expressed mural-associated genes, whereas the P3-low, R-like group expressed matrix-associated genes. Cross-checking against the original LuCA fine annotations^31^ showed that 97.9% of the L-like group comprised pericytes or smooth muscle cells and 98.0% of the R-like group comprised fibroblasts, with a small mesothelial group retained separately. Dancer preserved the expression differences between spatial L and R cells and made some of them more apparent. For example, MFAP5 showed only a modest difference between L and R in Raw, and its higher expression in R became clearer after correction, consistent with the reference. RGS5, MCAM, and MYLK remained more highly expressed in L cells. The reference expression profiles and original LuCA cell-type annotations thus supported the mural-like and matrix-associated identities of the spatial L and R populations, respectively.

Alveolar type 2 (AT2) cells provided a complementary setting in which to examine functional variation, as inflammatory and injury signals can activate stress-response programs accompanying transient repair-associated states^32, 33^. We therefore examined whether Dancer retained interpretable variation within this epithelial population. AT2 P3 and P5 programs showed similar activity gradients across the expression UMAP, with both increasing toward its lower part (Fig. 5i). Their high-loading genes were enriched for inflammatory responses, TNF–NFκB signaling and the unfolded protein response, with additional IL-6/JAK/STAT3 signaling in P3 and hypoxia-related expression in P5 (Fig. 5h).

Joint analysis of P3 and P5 activities identified J1 and J2 states in spatial AT2 cells and ref J1 and ref J2 states in the single-cell reference, with higher joint activity in both J2 groups (Fig. 5i, Methods). Within the measured panel, J1 and J2 showed clearer expression differences after Dancer correction than in Raw, and these differences more closely resembled those in the single-cell reference. J1 showed higher expression of the AT2 identity markers ABCA3, NKX2-1, and CLDN18. J2 showed higher expression of the response-associated genes CXCL2, ICAM1, and ATF3, as well as the protein-folding genes HSPA8 and DNAJA1 (Fig. 5j). Reference-guided panel expansion further predicted higher KRT8, KRT19, CLDN4 and SFN expression in J2, matching the direction of the reference-state differences and extending the interpretation to genes associated with transitional alveolar epithelial states in previous studies^33, 34^. Beyond individual markers, scores for published damage-associated transient progenitor (DATP)^35^ and alveolar differentiation intermediate (ADI)^33^ signatures were higher in J2 than J1 in both the reference and the expanded spatial profiles (Fig. 5j). Together, the measured expression differences, reference-guided predictions and whole-signature correspondence supported a repair- and transition-associated expression state at the high-P3/P5 end of AT2 variation.

Overall, these results show that Dancer scales to larger gene panels while reducing ectopic expression and preserving biologically interpretable within-cell-type heterogeneity.

## Discussion

In this work, we present Dancer, a reference-guided probabilistic framework that disentangles within-cell-type heterogeneity from transcript contamination in imaging-based ST data. Across Xenium datasets from breast, colorectal and lung cancers, Dancer improved expression specificity, and benchmark comparisons demonstrated superior overall decontamination performance relative to existing methods. Dancer also resolved spatially organized immune heterogeneity, tumor-associated expression gradients and biologically interpretable cell states. Three core features of Dancer’s design contribute to these results. First, by learning cell-type-specific expression programs from a single-cell reference assumed to be approximately free of transcript contamination and transferring them to the spatial model, Dancer uses the reference to guide the inference of intrinsic expression. These programs describe coordinated changes around a cell-type-specific baseline, with cell-specific activities capturing differences between states and continuous variation within them. At the same time, Dancer models observed spatial expression as separate contributions from intrinsic expression, neighboring-cell contamination, and background RNA, allowing program activities and source proportions to be inferred jointly. The inferred intrinsic component is then used to reconstruct decontaminated spatial expression profiles. Moreover, Dancer uses shared gene loadings for each cell type across the two modalities, so the separately inferred activities describe the same gene programs. This allows Dancer to match spatial cells to reference cells of the same type with similar program activities and impute the expression of unmeasured genes, expanding the spatial panel for more comprehensive functional interpretation.

Dancer also has its limitations that warrant further improvement. First, it requires a high-quality single-cell reference that is appropriate for the tissue and disease context and captures the relevant within-cell-type heterogeneity. Because Dancer learns its expression programs from this reference, missing cell states or biased expression profiles can lead to inaccurate decontamination and incomplete recovery of within-cell-type heterogeneity. Second, the spatial panels are required to include genes whose expression reflects the relevant within-cell-type variation. Even when the reference adequately represents this variation, insufficient coverage of these genes in the spatial panel can limit the reliable estimation of program activities, affecting the identification of within-cell-type variation and subsequent panel expansion based on activity matching. However, advances in spatial profiling, including Xenium 5K and the Atera whole-transcriptome platform, are expanding the range of genes that can be measured in situ^5, 12^. By providing direct measurements of more state-associated genes, these advances may help alleviate the constraints of limited panel coverage on heterogeneity analysis and decontamination.

Several directions for future development could enhance Dancer’s ability to resolve and interpret within-cell-type heterogeneity. First, program learning could incorporate reference genes beyond the shared panel. Jointly modeling correlated expression changes among genes within and beyond the spatial panel could improve the functional interpretation of programs and more closely connect program modeling with prediction of unmeasured genes. Additionally, future work could integrate reference samples from multiple patients with the same cancer type to learn programs representing a broader range of within-cell-type states. A broader reference could strengthen the biological interpretation of programs and provide a common basis for distinguishing intrinsic expression from contamination, supporting comparisons of cell states and their spatial organization across samples.

As imaging-based ST technologies continue to advance, distinguishing intrinsic expression variation from transcript contamination becomes increasingly important for understanding tissue biology. Dancer combines reference-guided expression program learning with explicit contamination modeling to generate decontaminated expression profiles while preserving within-cell-type heterogeneity. This framework supports a more faithful interpretation of cellular states and their relationships with the surrounding tissue microenvironment.

## Methods

### The model of Dancer

Dancer leverages a single-cell reference to guide the separation of intrinsic within-cell-type expression variation from transcript contamination in imaging-based spatial transcriptomics. This enables Dancer to remove contamination from spatial expression profiles while resolving heterogeneity among cells of the same type. Dancer assumes that the reference expression profiles are approximately free of transcript contamination and that, for each cell type, the reference captures the range of intrinsic expression variation present in the spatial sample. Dancer represents this variation through gene programs that capture coordinated expression changes across multiple genes, with the aim of improving the stability and functional interpretability of the inferred heterogeneity. For each cell type, Dancer learns gene program loadings from the single-cell reference and transfers them to the spatial model, where cell-specific program activities are inferred to reconstruct each spatial cell’s intrinsic expression. By combining reference-guided gene programs with a contamination-aware spatial model that simultaneously accounts for true cellular expression, contamination from neighboring cells, and background RNA, Dancer avoids misinterpreting contamination-driven expression patterns as genuine within-cell-type heterogeneity while recovering contamination-corrected expression profiles.

#### Inputs and notations

Let *G* denote the set of genes shared between the spatial panel and the single-cell reference, with *G* = |*G*| representing the number of genes in the shared panel. We denote the reference expression matrix by 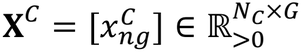 and the spatial count matrix by 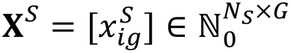, where *N*_*C*_ and *N*_*S*_ are the number of cells in the reference and the spatial sample, respectively. Reference cells are indexed by *n* = 1, …, *N*_*C*_, spatial cells by *i* = 1, …, *N*_*S*_, and genes by *g* = 1, …, *G*, with the same gene ordering in both matrices. The corresponding cell-type labels are 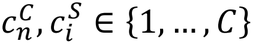, where *C* is the number of cell types, and *r*_*i*_ denotes the spatial coordinates of cell *i*. We define the cell-specific library sizes as 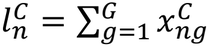 and 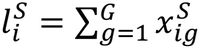 . For each cell type *c, K*_*C*_ denotes the number of gene programs used to represent within-type expression variation, determined using the selection procedure described below.

#### Reference-guided modeling of within-cell-type heterogeneity

Dancer assumes that the single-cell reference captures the within-cell-type expression variation present in the spatial data and is approximately free of transcript contamination. Under this assumption, Dancer learns cell-type-specific expression programs from the reference. We first establish a fixed expression baseline for each cell type. Let *T*_*C*_ denote the cells of type *c* in the reference. We normalize each cell by its panel-specific library size and average these expression compositions with equal weight:

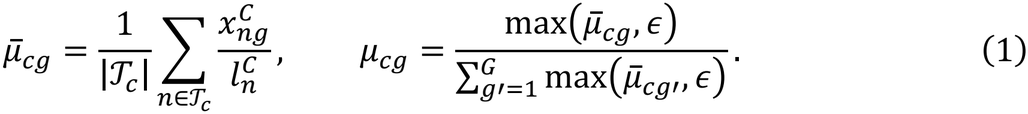

Here, ϵ > 0 is a small numerical constant that ensures positive baseline expression. The resulting vector *µ*_*c*_ sums to one and remains fixed throughout model fitting, providing a baseline expression for cells of the same type.

Following MacSGP^10^, we use a cell-type-specific loading matrix 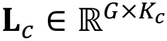 and cell-specific program activities 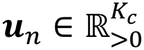 to model within-cell-type expression variation. Reference program activities are inferred using a cell-type-conditioned encoder with parameters *φ*_*C*_ . The encoder parameterizes a diagonal Gaussian variational posterior 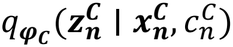 over the latent variable 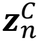 . Program activities are obtained by applying softplus elementwise, ensuring that they are nonnegative:

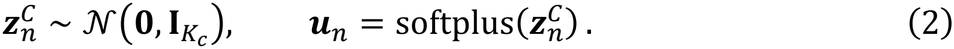

Therefore, multiple programs can be active simultaneously with activity levels varying across cells. The encoder architecture, latent priors and inference procedure are described in the model-fitting section below.

For a reference cell *n* of type *c*, we define its expression composition and Poisson observation model as

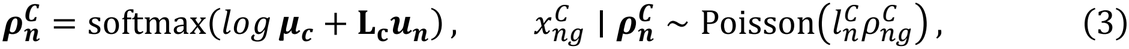

where softmax normalizes expression across genes. Equivalently,

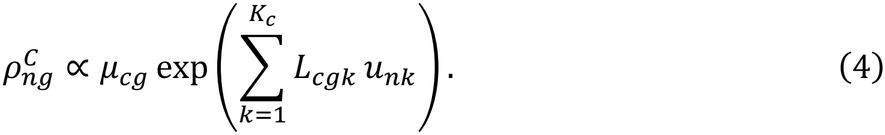

Thus, program activities modulate the reference baseline through coordinated multiplicative changes in relative gene expression. Each loading column can contain positive and negative values and is centered across genes, with 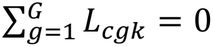. Cells of the same type share the baseline and loading matrix while differing in their program activities.

#### Contamination-aware model for spatial data

The observed counts for spatial cells are modeled as a mixture of contributions from the cell’s own expression, contamination from neighboring cells, and background RNA, with 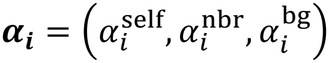 denoting their respective proportions in cell *i*. The source proportions are non-negative and sum to one, and their estimation and hierarchical prior are described in “Source proportions and spatial count likelihood” below. Let 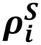 denote the cell’s intrinsic expression composition, *η*_*i*_ the composition of incoming neighbor contamination, and *ϕ* the background gene composition shared within a dataset. Their specific forms are described in the next three subsections, with each profile normalized across genes. The expected observed expression composition 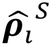 is

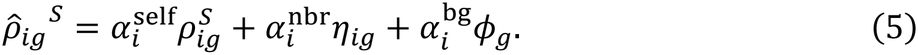

Multiplying this composition by the library size 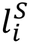 gives the expected observed gene counts 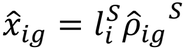.

#### Modeling intrinsic expression in spatial cells

For a spatial cell *i* of type *c*, a separate cell-type-conditioned encoder with parameters *φ*_*S*_ parameterizes the variational posterior 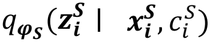 over its *K*_*C*_-dimensional latent state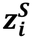. As in the reference model, this latent state is transformed into program activities *h*_*i*_, which modulate the reference-derived baseline *µ*_*c*_ through the shared loadings *L*_*c*_ to define the intrinsic expression composition 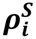:

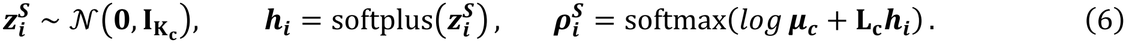

#### Modeling contamination from neighboring cells

Neighbor contamination is constructed from the intrinsic expression profiles of nearby cells. Let *N*(*i*) denote the set of the 20 nearest neighboring cells of cell *i*, with *ω* = (*ω*_*ij*_ ) _*j∈N(i)*_ denoting the relative contributions of neighboring cells to it. Here we assign a distance-dependent Dirichlet prior to *ω*_*i*_:

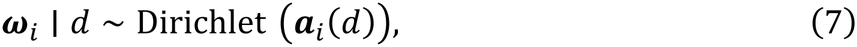

where *a*_*i*_(*d*) = (*a*_*ij*_(*d*))_*j∈N(i)*_, with

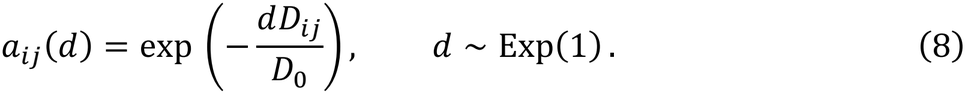

Here, 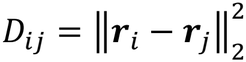 is the squared Euclidean distance between cells *i* and *j*, and *D*_0_ is the median, across all cells, of the squared distance to each cell’s sixth-nearest neighbor. This prior favors contributions from closer neighbors while allowing their relative weights to be inferred from the observed data.

Moreover, we define a positive gene-specific factor *γ*_*g*_, shared across spatial cells, to characterize the propensity of transcripts of gene *g* to diffuse from their cell of origin into neighboring cells, which is defined as follows:

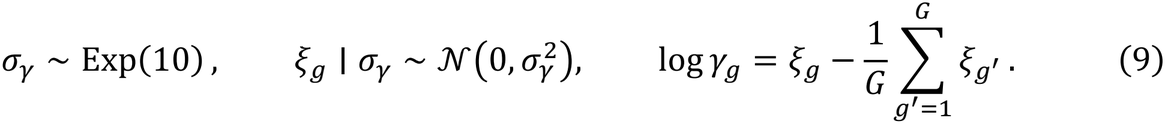

Then, the neighbor-derived contamination for cell *i* and gene *g*, denoted as *η*_*ig*_ is obtained as follows:

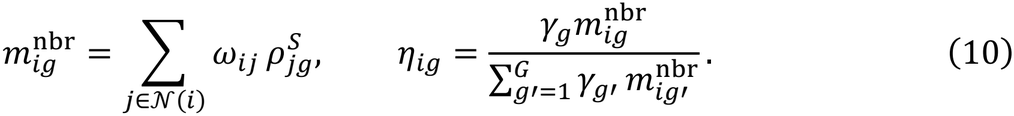

The resulting contamination profile adapts to the expression states of nearby cells and the diffusion propensity of each gene.

#### Modeling background expression

Within each dataset, background RNA is represented by a gene composition *ϕ* shared across spatial cells, with its contribution to each cell controlled by 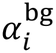. Dancer infers *ϕ* from spatial expression data under the following prior:

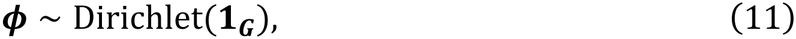

where *1*_*G*_ denotes a length-*G* vector of ones.

#### Source proportions and spatial count likelihood

A source-proportion encoder takes log-transformed spatial counts and produces three positive source scores, which are normalized to sum to one to obtain a point estimate of *α*_*i*_. These proportions are optimized jointly with the spatial program activities under the spatial observation model and the source-proportion prior. We model the source proportions with a global concentration vector *t* = (*t*_*q*_)_*q*∈{self,nbr,bg}_ as follows:

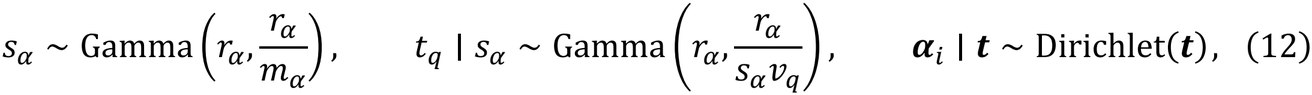

where *m* _*α*_ is fixed at 3, and both Gamma distributions use the shared fixed shape *r* _*α*_ = 10. The relative source weights *v* are defined as

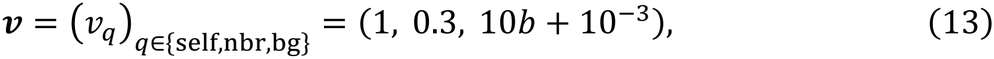

in which *b* is an empirical background scale estimated from the spatial training data. We first select the *⌊G*/50*⌋* shared genes with the lowest total counts across all spatial cells. For each cell, we divide the mean count across the selected genes by the mean count across all shared genes, then average these values across all cells to obtain *b*.

To model observed spatial counts, we first define the mean cell-derived intensity for gene *g* in cell *i* as:

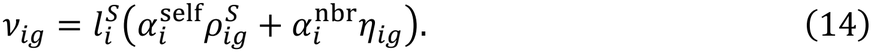

For each gene, we introduce an overdispersion parameter *δ*_*g*_, shared across spatial cells, and define the corresponding Gamma shape parameter 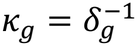. The latent cell-derived intensity Λ_*ig*_ is modeled as

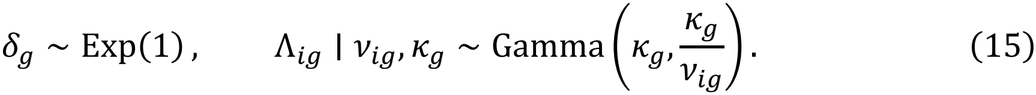

Finally, we incorporate the background intensity 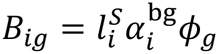 and model the observed spatial count as

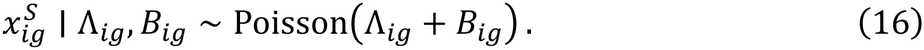

#### Parameter inference and model fitting

Overall, Dancer combines variational inference with maximum a posteriori (MAP) estimation. Variational inference is applied to the cell-specific latent states 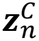 and 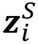, whose diagonal Gaussian posteriors are parameterized by the reference and spatial encoders, respectively. MAP estimation is used for

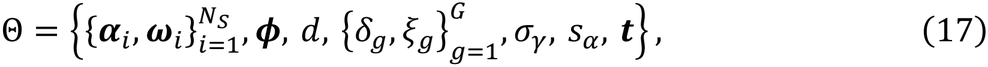

which contains parameters governing source proportions and neighbor contributions, together with their associated hyperparameters. The reference model is trained by maximizing the evidence lower bound (ELBO), whereas the spatial model is optimized using an ELBO augmented with the log-prior contributions of Θ.

Dancer first learns gene program loadings and trains the reference encoder using the single-cell reference. The learned program loadings are then held fixed when fitting the spatial model. Thus, the reference data determine the gene program loadings, whereas the spatial data inform cell-specific program activities and the contributions of intrinsic expression and other sources, which are inferred jointly. Both stages are optimized using Adam with minibatches, and the final model is selected based on the lowest spatial validation loss. Joint inference of spatial program activities and source contributions enables Dancer to disentangle intrinsic heterogeneity from transcript contamination.

#### Model outputs

After fitting, Dancer reconstructs intrinsic expression by setting *α*_*i*_ = (1,0,0) while retaining the observed panel-specific library size 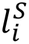. We draw 25 samples from each cell’s fitted latent posterior and generate corrected counts as follows:

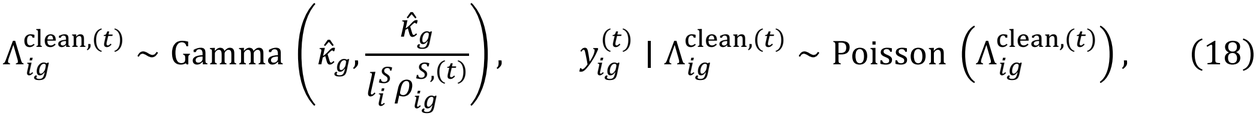

where 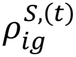 is the intrinsic expression composition obtained from the fitted latent posterior, 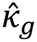 is the fitted Gamma shape parameter, and 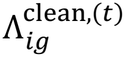 denotes the sampled intrinsic intensity. The corrected expression matrix is summarized by

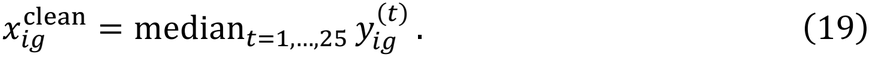

Dancer also exports the cell-type-specific loading matrices *L*_-_ and posterior mean program activities 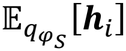 and 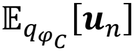 for spatial and reference cells, respectively.

#### Selection of the program numbers per cell type

We select *K*_*c*_ by fitting the reference model alone. Following MacSGP^10^, we start with one retained program per cell type and iteratively add one candidate program. In each round, cell types still undergoing selection are fitted jointly in the reference model. Previously learned program loadings are held fixed, while the new program loadings and reference encoder are trained. The initial model is trained for 50 epochs, followed by 25 additional epochs in each subsequent round. To evaluate each candidate program, we repeat the procedure with different random initializations, holding out a random subset of reference expression entries in each run. We assess predictive performance using the Poisson log-likelihood of held-out counts: ℓ(*x*; *m*) = *x* log *m* − *m* − log Γ (*x* + 1), where *x* is the observed count and *m* is its predicted mean. For each cell type, we compare the summed held-out log-likelihoods with the candidate program included or masked out. Let 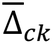 denote the mean improvement across runs and *p*_*ck*_ the *P* value from a one-sided, one-sample *t* test for positive improvement. We also evaluate program stability using *S*_*ck*_, defined as the mean pairwise absolute cosine similarity of the newly learned loading vectors across runs. A candidate program is retained if

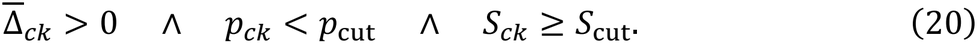

The default thresholds are *p*_cut_ = 0.10 and *S*_cut_ = 0.50. For each cell type, selection stops when a candidate first fails these criteria or when 10 programs have been retained, while other cell types continue independently. Once the program numbers are determined, we reinitialize and refit the model with the selected program numbers.

#### Model implementation

Dancer is implemented in Python using PyTorch for neural network computation and Pyro for probabilistic inference^36, 37^, with NumPy and SciPy for numerical and sparse-matrix operations^38, 39^. The five independent replicates used for program-number selection are parallelized across five NVIDIA GeForce RTX 4090 GPUs (24 GB each), with one replicate per GPU. Final model fitting is performed on a single GPU.

Each expression encoder consists of two hidden layers with 128 units, layer normalization, and a dropout rate of 0.05. The source-proportion encoder uses the same architecture without dropout. Models are optimized using Adam with a learning rate of 10^OP^ and batch sizes of 128 and 512 for reference and spatial data, respectively. By default, the spatial model is trained for 300 epochs without early stopping, and the checkpoint with the lowest spatial validation loss is selected. For validation, 10% of cells from each modality are held out using cell-type-stratified sampling.

For the breast cancer dataset, comprising 158,030 spatial cells, 26,031 reference cells and 306 shared genes, program-number selection required approximately 25 min of wall-clock time across five NVIDIA GeForce RTX 4090 GPUs. The final selected checkpoint was reached after approximately 1.2 h of model fitting on a single GPU.

### Data description and preprocessing

#### Breast cancer

We analyzed the human breast cancer Xenium dataset together with the matched Chromium single-cell FFPE reference and accompanying H&E and Visium measurements released by Janesick et al^3^. Published supervised Xenium annotations and the corresponding single-cell labels were retained. Cells with missing or unassigned labels and cells annotated as “Stromal & T Cell Hybrid” or “T Cell & Tumor Hybrid” were excluded. Cell-type names were harmonized across modalities, and shared gene-expression features with nonzero total expression in both modalities were retained. The resulting dataset contained 158,030 spatial cells, 26,031 reference cells, 17 cell types and 306 shared genes.

#### Colorectal cancer

We used the P2 colorectal cancer specimen profiled by Xenium, with Chromium Flex from the same specimen as the single-cell reference and matched Visium and Visium HD data for cross-platform validation^25^. Spatial cell-type labels were transferred from the reference using scANVI^40^. Spatial cells with at least 20 transcripts across the 407 shared genes were retained, yielding 317,294 cells across 24 cell types. Reference cells were required to have positive total expression across the shared panel, retaining all 34,790 reference cells.

#### Lung cancer

We analyzed the human lung cancer Xenium Prime 5K dataset using the LuCA lung cancer atlas as the single-cell reference^31^. Cell-type labels for spatial data were obtained using scANVI^40^. Filtering spatial cells for at least 20 transcripts across the 4,598 shared genes retained 271,371 cells across 22 cell types. The reference was constructed by cell-type-stratified sampling from LuCA, excluding cells labeled as “other” or “transitional club/AT2”, and comprised 151,970 cells and 17,764 genes. UMAP visualization of the uncorrected LuCA reference revealed separation associated with source datasets, suggesting batch effects. To reduce the risk of Dancer interpreting batch-associated variation as within-cell-type biological heterogeneity, we applied scVI using source dataset as the batch variable^41^. Denoised expression was averaged across batch conditions and rescaled by each cell’s original full-transcriptome total count. The shared genes were used for model fitting, and the full reference gene set was retained for panel expansion.

### Benchmarking analysis

Dancer was compared with uncorrected expression (Raw) and four decontamination methods: ResolVI^7^, DenoIST^11^, MisTIC^14^ and SPLIT^12^. The same cell-type annotations were used across all comparisons and provided to methods requiring them. Before evaluation, corrected expression matrices were aligned by cell and gene identifiers and restricted to the shared gene panel.

#### ResolVI

ResolVI separates intrinsic expression from neighborhood and background contamination using a probabilistic latent-variable model. We used 20 spatial neighbors and 10 latent dimensions, fitting the model in unsupervised mode for 100 epochs with a batch size of 512, a learning rate of 0.001 and a 20-epoch KL warmup. Corrected expression was obtained as the entrywise median of the corrected expression rates from 30 posterior samples.

#### DenoIST

DenoIST identifies contaminating counts using a Poisson mixture model informed by local transcript distributions. Version 1.1.0 was run with a neighborhood distance of 50 µm, 200 bins, a posterior cutoff of 0.6 and 10 initializations. The resulting filtered count matrix was used for evaluation.

#### MisTIC

MisTIC uses the local context of individual transcripts to infer and correct their cell assignments. We supplied the fixed cell-type labels and used the official neighborhood and correction-threshold settings, with 20 training epochs and *early_stop_pval=0*.*5*. Correction used the conservative mode for breast cancer and the *best* mode for colorectal cancer. Unnormalized corrected transcript counts were used for evaluation.

#### SPLIT

SPLIT purifies cell expression profiles using reference-guided estimates of cell-type mixtures. We ran SPLIT 0.3.0 with RCTD^42^ in doublet mode, using the same single-cell reference as Dancer. The minimum-count thresholds for both reference and spatial data were set to 10, with *UMI_min_sigma=100* and *CELL_MIN_INSTANCE=25*. Residual-contamination removal was disabled. Purified counts were used for evaluation, with original counts retained for input cells absent from the SPLIT output.

### Evaluation metrics

#### Residual contamination

We assessed residual contamination using the mutually exclusive co-expression rate (MECR) and positive marker purity (PMP)^11^. MECR quantifies the co-detection of marker genes specific to different cell types, which are expected not to be co-expressed within the same cell. For a gene *g*, let *A*_*g*_ contain cells with count-scale expression *x*_*ig*_ ≥ 0.5, and let *P* be the set of cross-type marker pairs with a nonempty union of expressing cells. We calculated

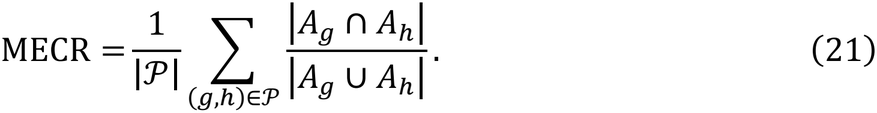

PMP measures the fraction of a cell’s expression contributed by markers of its annotated type. For marker set *M*_*c*_ and cells *I*_*c*_ of type *c*, we calculated

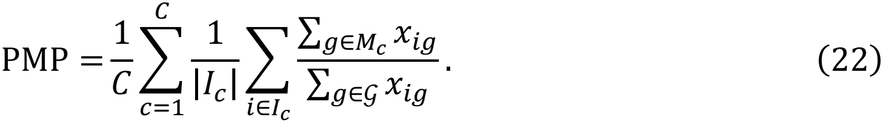

Cells with zero total expression contributed zero to PMP. Both metrics used all input cells; MECR was averaged equally across eligible gene pairs, and PMP was averaged within each cell type and then equally across types. Lower MECR indicates less cross-type marker co-expression, whereas higher PMP indicates a greater contribution from the cell’s own markers. Marker sets were derived from the single-cell reference and fixed across methods. Reference expression was normalized to 1,000 counts per cell and log-transformed, followed by one-versus-rest Wilcoxon tests in Scanpy^43^. Markers were required to have a Benjamini– Hochberg-adjusted *P* ≤ 0.05 and log fold change ≥ 0.5. The default detection thresholds were at least 40% of cells in the target type and at most 15% of all remaining cells, using count-scale expression ≥ 0.5 to define detection. PMP used all qualifying markers. For MECR, we retained the ten highest-ranked markers per type, removed genes assigned to more than one type and formed pairs between markers of different types.

#### Cell-type separability

We quantified cell-type separation using average silhouette width (ASW)^44^. For each decontamination method and the raw expression data (Raw), we retained cells with at least 20 total counts and restricted evaluation to cells shared across all methods and Raw. Each expression matrix was independently normalized to 10,000 counts per cell, log-transformed and reduced to 50 principal components. For cell *i*, let *a*_*i*_ denote its mean Euclidean distance to other cells of the same type, and let *b*_*i*_ denote the smallest mean distance to cells of any other type. The rescaled ASW is defined as

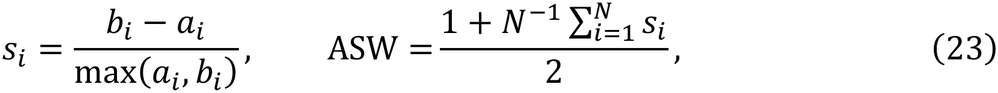

where *N* is the number of shared cells. We used *scib-metrics* for the calculation^45^. The resulting score ranges from 0 to 1, with higher values indicating clearer separation between cell types.

#### Reference concordance

Concordance with the single-cell references used for model fitting was assessed using representative-marker profiles in breast cancer (Fig. 2d, e) and cell-type expression profiles in colorectal cancer (Fig. 4b, bottom). Breast cancer dot plots summarized mean log-normalized expression and the fraction of cells with count-scale expression ≥0.5, with mean expression scaled by the maximum for each gene across the displayed methods and cell types. For the colorectal cancer analysis, NK cells were excluded because of their low abundance in Visium HD, with only 12 annotated bins, below the minimum threshold of 50. Counts were normalized to 10,000 per cell and averaged within each cell type. For each gene, the resulting mean expression values were further normalized to sum to one across cell types. Pearson correlations with Chromium were then calculated across genes for each cell type and averaged equally across cell types. Confidence intervals were estimated using 400 bootstrap resamples of genes.

#### Low-dimensional visualization

UMAP^46^ was used to visualize cell-type separability before and after decontamination. By default, cells with at least 20 total counts were retained. Expression across the shared panel was normalized to 10,000 counts per cell, log-transformed and reduced to 50 principal components using Scanpy^43^. A 20-nearest-neighbor graph was constructed, followed by two-dimensional UMAP with *min_dist=0*.*3* and random seed 42.

### Cross-platform validation of gene expression

#### Visium and Visium HD samples

In breast cancer, panel-expanded expression was compared with matched Visium measurements using the available Xenium–Visium registration^3^.

Xenium cells were assigned to their nearest Visium spot within the common tissue region, and predicted counts were summed per spot. Predicted and measured profiles were normalized to 10,000 counts over shared genes and log-transformed. Pearson correlations were calculated for each gene across corresponding spots containing assigned cells and positive total expression in both profiles. In colorectal cancer, registered Xenium cells, Visium spots and Visium HD bins were aggregated into a common 100 × 100 µm grid^25^. Counts were summed within each grid, normalized across the 407 shared genes, and log-transformed. ACTA2 expression from Raw and Dancer was separately correlated with that from Visium and Visium HD using the same grids occupied by both platforms with positive total counts. Pearson and Spearman correlations, along with their two-sided *P* values, were calculated using SciPy^39^.

#### H&E image

The post-Xenium hematoxylin and eosin (H&E) image and accompanying registration information were obtained from the 10x Genomics breast cancer dataset described by Janesick et al^3^. The H&E image was aligned with Xenium coordinates using the provided transformation to examine ACTA2 expression in the anatomical context of a duct. We compared ACTA2 expression between annotated myoepithelial cells and adjacent epithelial or tumor cells using Raw and Dancer. Cell-type discrimination was quantified using the area under the receiver operating characteristic curve (AUROC) and Spearman correlation between log-normalized ACTA2 expression and binary myoepithelial labels. Image scale bars were determined from the recorded pixel size and spatial registration.

### Lineage-specific co-expression analysis

In lung cancer, we adapted the co-expression evaluation strategy used in ResolVI^7^ to assess whether decontamination reduced cross-lineage co-expression while preserving reference-supported co-expression. Lineage markers were selected from predefined identity-gene sets using differential expression and specificity in the scVI-processed reference, after setting count-scale values below 0.5 to zero. For each gene and expression matrix, a two-component Poisson mixture, adapted from the ResolVI analysis code, was fitted to expression normalized to 10,000 counts per cell. Cells assigned to the higher-expression component by posterior probability were considered positive. Gene-pair co-expression was measured by the Jaccard index: the number of cells positive for both genes divided by the number positive for either gene. The reference defined two fixed pair sets: markers from the same lineage with co-expression ≥0.10 and markers from different lineages with co-expression ≤0.05. Raw and Dancer were compared using these same pairs. Changes in co-expression and in absolute deviation from the reference were assessed within each pair set using two-sided paired Wilcoxon signed-rank tests^47^, with Benjamini–Hochberg correction^48^ across the four tests. Each gene pair constituted one paired observation.

### Identification of within-cell-type states

#### Expression-based clustering

In breast cancer, B cells were partitioned into B1 and B2 by applying DBSCAN^49^ to their coordinates in the Dancer expression UMAP. In colorectal cancer, macrophages were analyzed on a fixed random sample of 12,000 cells with positive total expression in both Raw and Dancer. Expression was normalized to 10,000 counts per cell, log-transformed and standardized gene by gene, followed by PCA, nearest-neighbor graph construction and Leiden clustering^43, 50^. The five initial Dancer clusters were merged according to marker expression into SPP1, CD163 and MMP12 states. In lung cancer, stromal cells were analyzed using the same expression-processing framework, with Leiden resolution chosen to recover two major spatial populations, labeled L and R. Reference stromal cells were clustered independently, and the principal populations were matched to spatial states using identity-gene expression and the original LuCA fine annotations. A small mesothelial reference population was retained separately.

#### Program-based clustering

In breast cancer, CD8 T cells neighboring exclusively B1 or B2 cells were clustered using standardized P3 and P4 activities. A nearest-neighbor graph was constructed in this activity space, and Leiden resolution was selected using silhouette scores subject to a minimum cluster-size requirement. The resulting states were displayed on an independently computed expression UMAP. In lung cancer, AT2 cells were grouped separately in spatial and reference data using P3 and P5 activities. Activities were log-transformed with *log1p*, standardized within each modality and partitioned by two-cluster K-means in scikit-learn^51^. Groups were designated J1 and J2 in order of increasing mean standardized activity. State labels were fixed for subsequent gene-expression, spatial and panel-expansion analyses.

### Differential expression and state-associated gene expression

Differential expression between cell states was assessed using two-sided Wilcoxon rank-sum tests in Scanpy^43^, with tie correction and Benjamini–Hochberg adjustment across tested genes^48^. Counts were normalized to 10,000 per cell over the full analysis panel and transformed with *log1p*. In breast cancer, B2 was compared with B1 using Dancer expression; volcano plots displayed Scanpy-derived log2 fold changes against −log10 adjusted *P* values. State labels were held fixed for subsequent expression comparisons, and representative genes were selected using their biological functions and observed expression patterns. For breast CD8 T-cell heatmaps, expression was standardized gene by gene across the analyzed cells before averaging within each state. Lung stromal comparisons showed state-specific mean log-normalized expression in Raw, Dancer and the reference. For lung AT2 dot plots, gene-wise standardization was performed separately within each modality before calculating state means; dot size represented the fraction of cells with count-scale expression ≥0.5.

### Gene set enrichment analysis

#### Loading-based over-representation analysis

Following the program-interpretation approach used in MacSGP^10^, we annotated selected programs in breast, colorectal and lung cancer by ranking genes according to their cell-type-specific loadings. The 100 highest- and 100 lowest-loading genes were tested separately against the MSigDB Hallmark collection^20^ using Enrichr^52^ and its default background. Enrichment was assessed by one-sided Fisher’s exact tests, using the returned Benjamini–Hochberg-adjusted *P* values across the tested collection for each loading end. Functionally relevant terms with adjusted *P* ≤ 0.05 were displayed. Bidirectional bars distinguished the high- and low-loading ends, with lengths representing −log10 unadjusted *P* values.

#### Expression–activity-ranked gene set enrichment analysis

In breast cancer, we compared functional associations of B-cell and CD8 T-cell programs between the measured and expanded panels. For each panel, genes were ranked by Spearman correlation between their log-normalized expression and program activity, using the same cells and fixed activities. Constant genes were excluded; gene symbols were mapped to Entrez identifiers, retaining the largest absolute correlation for duplicate mappings. Preranked gene set enrichment analysis (GSEA)^23^ was performed with clusterProfiler^53^ using the adaptive multilevel fgsea algorithm^54^ for GO Biological Process, human KEGG and Hallmark gene sets, retaining sets with 10–500 genes represented in the ranking. Normalized enrichment scores (NES) indicated enrichment toward genes positively or negatively correlated with activity. *P* values were adjusted by the Benjamini–Hochberg procedure within each collection and ranking. Displayed significance thresholds were adjusted *P* ≤ 0.05 for GO and Hallmark and *P* ≤ 0.10 for KEGG. Bubble plots showed NES, adjusted *P* values and the number of leading-edge genes contributing to enrichment.

### Validating spatial patterns of inferred program activity

We compared Dancer program activities with gene-specific expression patterns inferred by MMM, using the authors’ MCube R implementation^19^. Analyses included the 4,862 breast B cells and a fixed, spatially stratified subset of 4,000 lung stromal cells. Inputs comprised Raw counts, cell-centroid coordinates, shared-panel library sizes and cell-type expression profiles from the corresponding single-cell reference. Each position represented one annotated cell, with the target cell-type proportion fixed at one. MMM was fitted by penalized quasi-likelihood, and spatial variation was assessed using its kernel-based score tests, with kernel-specific *P* values combined by ACAT and Benjamini–Hochberg correction across tested genes. We extracted the fitted gene-specific expression-variation estimates at each cell and compared their tissue distributions with Dancer activity at the same cell identifiers. Agreement was quantified by Spearman correlation with two-sided *P* values, focusing on BANK1 and B-cell P3 in breast cancer and MYLK and stromal P3 in lung cancer.

### Spatial neighborhood analysis

Spatial neighborhoods were defined using the original cell-centroid coordinates and annotations. For breast B cells and lung stromal cells, we identified all other annotated cells within 30 µm, calculated the fraction of each neighboring cell type and averaged these fractions within each focal state. Cells without neighbors were omitted from these averages. Breast CD8 T cells were assigned to B1-associated, B2-associated, shared or other neighborhoods according to whether B1 and B2 cells occurred within this radius. State-composition comparisons used CD8 cells associated exclusively with B1 or B2, with proportions calculated within each neighborhood group. In colorectal cancer, we assessed expression similarity to nearby cells from different lineages (Fig. 4b, top). Fine annotations were grouped into six broad lineages: tumor, T/NK, myeloid, stromal/endothelial, B/plasma and epithelial. For each cell, we selected its 15 nearest other cells and retained neighbors from different lineages. Count vectors were L2-normalized, and the mean neighbor vector was normalized again before calculating cosine similarity with the focal cell. The same neighbor sets were used for Raw and Dancer. Similarities were averaged by focal-cell lineage; cells without eligible neighbors were excluded, and zero-expression vectors contributed zero similarity.

### Gene expression and program activity in relation to tumor proximity

In colorectal cancer, tumor proximity was quantified using local cell composition or spatial distance. For SPP1 macrophages, we calculated the tumor-cell fraction among the 30 nearest other cells. For CD163 macrophages, distance to the tumor edge was estimated by finding the nearest tumor-cell centroid, subtracting the two cells’ area-equivalent radii, and truncating negative values to zero. For goblet cells, tumor regions were defined on a 20-µm grid from Gaussian-smoothed cell densities (σ=40 µm), requiring a tumor-cell fraction ≥0.5 and retaining connected regions containing at least 100 tumor cells. Distances to these regions were calculated by a Euclidean distance transform, with cells inside a region assigned zero distance; goblet cells beyond the 95th distance percentile were excluded from trend analysis. Associations with program activity and log-normalized gene expression were assessed using per-cell Spearman correlations and summarized in equal-frequency bins, with zero edge distances grouped separately for CD163 cells. Where shown, error bars represented the binned mean ± 1.96 times the standard error of the mean. Goblet program–distance associations were additionally fitted by ordinary least squares with 95% confidence bands. In lung cancer, stromal cells within 20 µm of the estimated tumor edge and those at least 80 µm away were designated near and far, respectively (Fig. 5d), using the same edge-distance definition as for CD163 cells. Raw and Dancer were compared on identical cells and fixed proximity groups. Near-versus-far differential expression was tested separately in each matrix using Scanpy Wilcoxon tests with tie correction and Benjamini–Hochberg adjustment^43, 48^. Signed Wilcoxon Z statistics were compared across methods, with positive values indicating higher expression near the tumor.

### Cell–cell communication analysis

Cell–cell communication in breast cancer was inferred separately from Raw, Dancer and Chromium expression using CellChat 2.2.0^15^ and its human ligand–receptor database (Fig. 3b). Each analysis used the same 306-gene panel, fixed cell-type annotations and log-normalized counts. After identifying overexpressed signaling genes and interactions, communication probabilities were estimated using trimean expression summaries within cell types. Significance was assessed by randomly permuting cell-type labels 100 times and recalculating communication probabilities. A directed communication edge was considered significant at *P* < 0.05, with at least ten cells in both the sending and receiving groups. We compared the CellChat results from Raw and Dancer separately with those from the matched Chromium reference. At the edge level, a match required the same sending cell type, receiving cell type and ligand–receptor pair. At the pair level, a ligand–receptor pair was counted once per analysis if it appeared in at least one significant directed edge. At each level, precision was the proportion of edges or pairs identified in the spatial analysis that were also identified in Chromium, and recall was the proportion identified in Chromium that were also identified in the spatial analysis. In the ligand-receptor network plots, nodes represent cell types arranged in the same circular layout, arrows show significant interactions from sender to receiver, and arrow widths reflect CellChat communication probabilities on a shared scale across the three analyses.

### REFERENCE-guided gene panel expansion

To estimate expression beyond the measured spatial panel, we matched each spatial cell to reference cells of the same type using the learned program activities. Pearson similarity between the spatial activity vector *h*_*i*_ and reference activity vector *u*_*n*_ was calculated across all retained programs for that cell type. The ten most similar reference cells were selected, and their count-scale expression values were averaged with equal weights to predict each gene’s expression. For cell types with only one or two retained programs, matching used Euclidean distance between activity vectors. Model parameters, cell-type labels and state assignments were held fixed during expansion.

### State-signature scoring

We assessed correspondence between lung AT2 states and published repair-associated expression programs using fixed damage-associated transient progenitor (DATP) and alveolar differentiation intermediate (ADI) gene sets. The human DATP signature was obtained from the author-released resources of Melms et al^35^. Krt8-positive ADI markers were obtained from the supplementary data of Strunz et al.^33^ and converted to human genes through one-to-one orthologue mappings from Mouse Genome Informatics. All signature genes represented in the full reference panel were retained, yielding 147 DATP and 353 ADI genes. Reference and panel-expanded spatial expression were separately normalized to 10,000 counts per cell and log-transformed. Scores were calculated with Scanpy’s *score_genes*^43^, following the expression-matched control approach of Tirosh et al^55^. J2–J1 differences were summarized within each modality by Cohen’s *d*, defined as the difference in mean scores divided by the pooled within-group standard deviation; positive values indicated higher scores in J2.

## Data availability

### Spatial datasets

Three Xenium datasets were included in this study: human breast cancer (167,780 cells, 313-gene panel; https://www.10xgenomics.com/products/xenium-in-situ/preview-dataset-human-breast), human colorectal cancer (340,837 cells, 422-gene panel, sample P2; https://www.10xgenomics.com/platforms/visium/product-family/dataset-human-crc), and human lung cancer (278,328 cells, 5,001-gene Xenium Prime 5K panel; https://www.10xgenomics.com/datasets/xenium-human-lung-cancer-post-xenium-technote).

Matched Visium and Visium HD measurements from the colorectal cancer specimen were used for cross-platform comparisons.

### Single-cell references

Each spatial dataset was analyzed with a labeled single-cell reference. The breast cancer section used the matched Chromium single-cell FFPE dataset released with the same study^3^. The colorectal cancer section used the Chromium Flex dataset from the same P2 specimen, available from the colorectal cancer dataset collection listed above. The lung cancer section used the public LuCA lung cancer atlas^31^, available through CELLxGENE (https://cellxgene.cziscience.com/collections/edb893ee-4066-4128-9aec-5eb2b03f8287).

Model fitting was performed on genes shared between each spatial panel and its corresponding reference; the broader reference gene sets were retained for panel expansion.

## Code availability

Dancer model and related tutorials are available at https://github.com/YangLabHKUST/Dancer.

## Acknowledgements

This work was supported by the Young Student Basic Research Program under the National Natural Science Foundation of China (125B2027).

## Author contributions

Y. L. conceived the study and supervised the project. Z. L. designed, implemented, and validated Dancer. Y. C. and C. Y. contributed to the experimental design and helped with analyzing the results. Z. L. and Y. L. wrote the manuscript with input from all the authors.

## References

1. Chen, K. H., Boettiger, A. N., Moffitt, J. R., Wang, S. & Zhuang, X. Spatially resolved, highly multiplexed RNA profiling in single cells. Science 348, aaa6090 10.1126/science.aaa6090 (2015).

2. Eng, C.-H. L., et al. Transcriptome-scale super-resolved imaging in tissues by RNA seqFISH+. Nature 568, 235–239 10.1038/s41586-019-1049-y (2019).

3. Janesick, A., et al. High resolution mapping of the tumor microenvironment using integrated single-cell, spatial and in situ analysis. Nat Commun 14, 8353 10.1038/s41467-023-43458-x (2023).

4. He, S., et al. High-plex imaging of RNA and proteins at subcellular resolution in fixed tissue by spatial molecular imaging. Nat Biotechnol 40, 1794–1806 10.1038/s41587-022-01483-z (2022).

5. Wu, S., et al. Resolving the immune response across clonal cancer evolution in situ with Atera whole transcriptome profiling. Preprint at 10.64898/2026.08.03.741871 (2026)

6. Petukhov, V., et al. Cell segmentation in imaging-based spatial transcriptomics. Nat Biotechnol 40, 345–354 10.1038/s41587-021-01044-w (2022).

7. Ergen, C. & Yosef, N. ResolVI: addressing noise and bias in spatial transcriptomics. Nat Methods 23, 2003–2014 10.1038/s41592-026-03212-9 (2026).

8. Tiesmeyer, S., et al. Identifying 3D signal overlaps in spatial transcriptomics data with ovrlpy. Nat Biotechnol, 10.1038/s41587-026-03004-8 (2026).

9. Mitchel, J., Gao, T., Petukhov, V., Cole, E. & Kharchenko, P. V. Impact and correction of segmentation errors in spatial transcriptomics. Nat Genet 58, 434–444 10.1038/s41588-025-02497-4 (2026).

10. Zeng, Y., et al. Mapping cell-type-specific spatial gene programs uncovers tissue architecture and microenvironment organization. Nat Commun 17, 11610 10.1038/s41467-026-78447-3 (2026).

11. Kwok, A. W. C., et al. Denoising image-based spatial transcriptomics data with DenoIST. Preprint at 10.1101/2025.11.13.688387 (2025)

12. Bilous, M., et al. Resolving sensitivity, specificity and signal contamination in Xenium spatial transcriptomics. Nat Methods 23, 1152–1162 10.1038/s41592-026-03089-8 (2026).

13. Chen, Y., et al. Accurate and scalable decontamination of imaging-based spatial transcriptomics via optimal transport. Preprint at 10.64898/2026.09.09.750350 (2026)

14. Yang, Y., et al. MisTIC: missegmented transcript inference correction for improved spatial transcriptomics analysis. Preprint at 10.64898/2025.12.11.693759 (2025)

15. Jin, S., et al. Inference and analysis of cell-cell communication using CellChat. Nat Commun 12, 1088 10.1038/s41467-021-21246-9 (2021).

16. Yokoyama, K., et al. BANK regulates BCR-induced calcium mobilization by promoting tyrosine phosphorylation of IP3 receptor. EMBO J 21, 83–92 10.1093/emboj/21.1.83 (2002).

17. Rosenbaum, M., et al. MZB1 is a GRP94 cochaperone that enables proper immunoglobulin heavy chain biosynthesis upon ER stress. Genes Dev 28, 1165–1178 10.1101/gad.240762.114 (2014).

18. Bilska, A., et al. Immunoglobulin expression and the humoral immune response is regulated by the non-canonical poly(A) polymerase TENT5C. Nat Commun 11, 2032 10.1038/s41467-020-15835-3 (2020).

19. Wang, Z., et al. A unified framework for identification of cell-type-specific spatially variable genes in spatial transcriptomic studies. Proc Natl Acad Sci USA 122, e2503952122 10.1073/pnas.2503952122 (2025).

20. Liberzon, A., et al. The Molecular Signatures Database hallmark gene set collection. Cell Syst 1, 417–425 10.1016/j.cels.2015.12.004 (2015).

21. Noël, G., et al. Functional Th1-oriented T follicular helper cells that infiltrate human breast cancer promote effective adaptive immunity. J Clin Invest 131, e139905 10.1172/JCI139905 (2021).

22. Sallusto, F., Lenig, D., Förster, R., Lipp, M. & Lanzavecchia, A. Two subsets of memory T lymphocytes with distinct homing potentials and effector functions. Nature 401, 708–712 10.1038/44385 (1999).

23. Subramanian, A., et al. Gene set enrichment analysis: a knowledge-based approach for interpreting genome-wide expression profiles. Proc Natl Acad Sci USA 102, 15545–15550 10.1073/pnas.0506580102 (2005).

24. Li, S., et al. An integrated map of fibroblastic populations in human colon mucosa and cancer tissues. Commun Biol 5, 1326 10.1038/s42003-022-04298-5 (2022).

25. Oliveira, M. F. d., et al. High-definition spatial transcriptomic profiling of immune cell populations in colorectal cancer. Nat Genet 57, 1512–1523 10.1038/s41588-025-02193-3 (2025).

26. Zhang, L., et al. Single-cell analyses inform mechanisms of myeloid-targeted therapies in colon cancer. Cell 181, 442–459.e429 10.1016/j.cell.2020.03.048 (2020).

27. Matusiak, M., et al. Spatially segregated macrophage populations predict distinct outcomes in colon cancer. Cancer Discov 14, 1418–1439 10.1158/2159-8290.CD-23-1300 (2024).

28. Qi, J., et al. Single-cell and spatial analysis reveal interaction of FAP+ fibroblasts and SPP1+ macrophages in colorectal cancer. Nat Commun 13, 1742 10.1038/s41467-022-29366-6 (2022).

29. Renner, M., et al. DMBT1 confers mucosal protection in vivo and a deletion variant is associated with Crohn’s disease. Gastroenterology 133, 1499–1509 10.1053/j.gastro.2007.08.007 (2007).

30. Travaglini, K. J., et al. A molecular cell atlas of the human lung from single-cell RNA sequencing. Nature 587, 619–625 10.1038/s41586-020-2922-4 (2020).

31. Salcher, S., et al. High-resolution single-cell atlas reveals diversity and plasticity of tissue-resident neutrophils in non-small cell lung cancer. Cancer Cell 40, 1503–1520.e1508 10.1016/j.ccell.2022.10.008 (2022).

32. Choi, J., et al. Inflammatory signals induce AT2 cell-derived damage-associated transient progenitors that mediate alveolar regeneration. Cell Stem Cell 27, 366–382.e367 10.1016/j.stem.2020.06.020 (2020).

33. Strunz, M., et al. Alveolar regeneration through a Krt8+ transitional stem cell state that persists in human lung fibrosis. Nat Commun 11, 3559 10.1038/s41467-020-17358-3 (2020).

34. Kobayashi, Y., et al. Persistence of a regeneration-associated, transitional alveolar epithelial cell state in pulmonary fibrosis. Nat Cell Biol 22, 934–946 10.1038/s41556-020-0542-8 (2020).

35. Melms, J. C., et al. A molecular single-cell lung atlas of lethal COVID-19. Nature 595, 114–119 10.1038/s41586-021-03569-1 (2021).

36. Paszke, A., et al. PyTorch: an imperative style, high-performance deep learning library. In: Advances in Neural Information Processing Systems. Curran Associates, Inc. (2019).

37. Bingham, E., et al. Pyro: deep universal probabilistic programming. J Mach Learn Res 20, 1–6 (2019).

38. Harris, C. R., et al. Array programming with NumPy. Nature 585, 357–362 10.1038/s41586-020-2649-2 (2020).

39. Virtanen, P., et al. SciPy 1.0: fundamental algorithms for scientific computing in Python. Nat Methods 17, 261–272 10.1038/s41592-019-0686-2 (2020).

40. Xu, C., et al. Probabilistic harmonization and annotation of single-cell transcriptomics data with deep generative models. Mol Syst Biol 17, e9620 10.15252/msb.20209620 (2021).

41. Lopez, R., Regier, J., Cole, M. B., Jordan, M. I. & Yosef, N. Deep generative modeling for single-cell transcriptomics. Nat Methods 15, 1053–1058 10.1038/s41592-018-0229-2 (2018).

42. Cable, D. M., et al. Robust decomposition of cell type mixtures in spatial transcriptomics. Nat Biotechnol 40, 517–526 10.1038/s41587-021-00830-w (2022).

43. Wolf, F. A., Angerer, P. & Theis, F. J. SCANPY: large-scale single-cell gene expression data analysis. Genome Biol 19, 15 10.1186/s13059-017-1382-0 (2018).

44. Rousseeuw, P. J. Silhouettes: a graphical aid to the interpretation and validation of cluster analysis. J Comput Appl Math 20, 53–65 10.1016/0377-0427(87)90125-7 (1987).

45. Luecken, M. D., et al. Benchmarking atlas-level data integration in single-cell genomics. Nat Methods 19, 41–50 10.1038/s41592-021-01336-8 (2022).

46. McInnes, L., Healy, J. & Melville, J. UMAP: Uniform Manifold Approximation and Projection for Dimension Reduction. Preprint at 10.48550/arXiv.1802.03426 (2018)

47. Wilcoxon, F. Individual comparisons by ranking methods. Biometrics Bull 1, 80–83 10.2307/3001968 (1945).

48. Benjamini, Y. & Hochberg, Y. Controlling the false discovery rate: a practical and powerful approach to multiple testing. J R Stat Soc B 57, 289–300 10.1111/j.2517-6161.1995.tb02031.x (1995).

49. Ester, M., Kriegel, H.-P., Sander, J. & Xu, X. A density-based algorithm for discovering clusters in large spatial databases with noise. In: Proceedings of the Second International Conference on Knowledge Discovery and Data Mining. AAAI Press (1996).

50. Traag, V. A., Waltman, L. & van Eck, N. J. From Louvain to Leiden: guaranteeing well-connected communities. Sci Rep 9, 5233 10.1038/s41598-019-41695-z (2019).

51. Pedregosa, F., et al. Scikit-learn: machine learning in Python. J Mach Learn Res 12, 2825–2830 (2011).

52. Kuleshov, M. V., et al. Enrichr: a comprehensive gene set enrichment analysis web server 2016 update. Nucleic Acids Res 44, W90–W97 10.1093/nar/gkw377 (2016).

53. Wu, T., et al. clusterProfiler 4.0: a universal enrichment tool for interpreting omics data. Innovation 2, 100141 10.1016/j.xinn.2021.100141 (2021).

54. Korotkevich, G., et al. Fast gene set enrichment analysis. Preprint at 10.1101/060012 https://www.biorxiv.org/content/10.1101/060012v3 (2021)

55. Tirosh, I., et al. Dissecting the multicellular ecosystem of metastatic melanoma by single-cell RNA-seq. Science 352, 189–196 10.1126/science.aad0501 (2016).

